# Integrin α6β4 recognition of a linear motif of bullous pemphigoid antigen BP230 controls its recruitment to hemidesmosomes

**DOI:** 10.1101/402123

**Authors:** José A Manso, Maria Gómez-Hernández, Arturo Carabias, Noelia Alonso-García, Inés García-Rubio, Maaike Kreft, Arnoud Sonnenberg, José M de Pereda

**Affiliations:** Instituto de Biología Molecular y Celular del Cáncer, Consejo Superior de Investigaciones Científicas - University of Salamanca, Campus Unamuno, 37007 Salamanca, Spain.; Centro Universitario de la Defensa, ctra. Huesca s/n, 50090 Zaragoza, Spain.; Netherlands Cancer Institute, Plesmanlaan 121, 1066 CX Amsterdam, The Netherlands.

**Keywords:** Cell adhesion, epithelia, plakins, proteinprotein interactions

## Abstract

Mechanical stability of epithelia requires firm attachment to the basement membrane via hemidesmosomes. Dysfunction of hemidesmosomal proteins causes severe skin blistering diseases. Two plakins, plectin and BP230 (BPAG1e), link the integrin α6β4 to intermediate filaments in epidermal hemidesmosomes. Here, we show that a linear sequence within the isoform-specific N-terminal region of BP230 binds to the third and fourth FnIII domains of β4. The crystal structure of the complex and mutagenesis analysis revealed that BP230 binds between the two domains of β4. BP230 induces closing of the two FnIII domains that are looked in place by an inter-domain ionic clasp required for binding. Disruption of the BP230-β4 interface prevents the recruitment of BP230 to hemidesmosomes in human keratinocytes, revealing a key role of the BP230-β4 interaction for hemidesmosome assembly. Phosphomimetic substitutions in β4 and BP230 disrupt binding. Our study provides insights into the molecular mechanisms of hemidesmosome architecture and regulation.

## Introduction

Hemidesmosomes (HDs) are junctional complexes that mediate the firm attachment of epithelial cells to the basement membrane to maintain the integrity of epithelial tissues (Walko et al., 2015). Pseudostratified and stratified epithelia, such as the epidermis, assemble classic type I HDs. These are rivet-like structures that contain three transmembrane proteins, the integrin α6β4, the bullous pemphigoid antigen BP180 (also known as BPAG2 or collagen XVII), and the tetraspanin CD151; and two intracellular proteins of the plakin family, plectin and BP230 (also known as BPAG1e). α6β4 and BP180 bind to laminin-332 in the epidermal basement membrane (Van den Bergh et al., 2011; Wilhelmsen et al., 2006), and are connected to the keratin intermediate filaments via plectin and BP230 (Geerts et al., 1999; Niessen et al., 1997a; Rezniczek et al., 1998), thereby linking the extracellular matrix to the cytoskeleton. Simple epithelia, such as that in the intestine, assemble type II HDs that contain α6β4 and plectin but lack BP180 and BP230.

The integrin α6β4 is an essential component of HDs and a hub of the HD protein-interaction network. Most of the intracellular interactions of α6β4 are established by the β4 subunit. The β4 cytodomain (∼1000 residues) is unique in the integrin family, it contains a Calxβ and four fibronectin type-III domains arranged in two pairs (FnIII-1,2 and FnIII-3,4) separated by a region named the connecting segment (CS); a C-terminal tail (C-tail) extends downstream of the FnIII-4 (Figure 1A). The FnIII-1,2 and the beginning of the CS bind to the actin-binding domain (ABD) of plectin (de Pereda et al., 2009; Geerts et al., 1999). The final part of the CS and the C-tail make a second site of contact with the plakin domain of plectin (Koster et al., 2004; Rezniczek et al., 1998). The FnIII-3,4 and part of the CS bind to BP230 (Hopkinson and Jones, 2000; Koster et al., 2003). The FnIII-3 also interacts with the cytoplasmic domain of BP180 (Koster et al., 2003). Recently, the region FnIII-3,4-C-tail has been shown to interact with Solo (ARHGEF40), a guanine nucleotide exchange factor of the RhoA small GTPase, which is required for the formation of HDs in mammary epithelial cells (Fujiwara et al., 2018).

**Figure 1.**
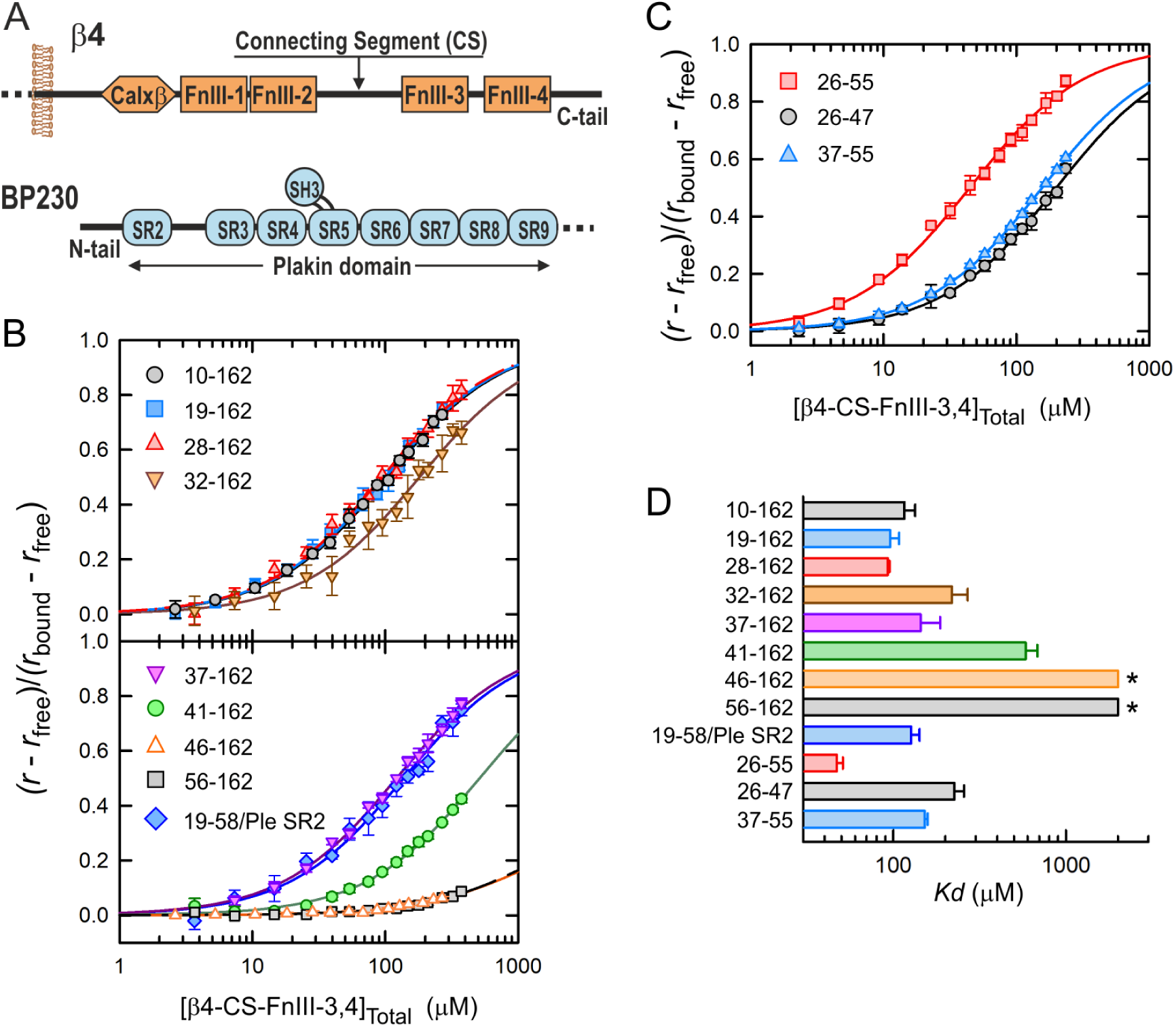
Mapping the β4-binding site in BP230. **(A)** Domain organization of the cytoplasmic region of β4 and the N-terminal region of BP230. (**B**) Equilibrium binding of β4-CS-FnIII-3,4 to Oregon Green-labeled fragments of BP230, measured by fluorescence anisotropy (r). Data are presented as fractional saturation. Lines are the fit of a one-to-one binding model. (**C**) Equilibrium binding of β4-CS-FnIII-3,4 to fluorescein-labeled peptides of the N-tail of BP230; data and the fits are shown as in B. (**D**) Bar-chart of the *Kd* of the interaction between β4-CS-FnIII-3,4 and the BP230 fragments. Error bars show the standard deviation (SD) from 2-4 experiments. Asterisks indicate minimal values compatible with the data.

BP230 is the epidermal-specific splice variant coded by the BPAG1/dystonin gene *DST* (Leung et al., 2001). BP230 has the characteristic three-segment structure shared by classic plakins. The N-terminal region contains a short isoform-specific N-terminal tail (N-tail) followed by a plakin domain (∼1000 residues) that consists of eight spectrin repeats (SR2 to SR9) and an SH3 domain (Jefferson et al., 2007; Sonnenberg et al., 2007) (Figure 1A). This region interacts with α6β4 and with BP180 (Koster et al., 2003). The central region is occupied by a coiled-coil rod domain that acts as a spacer (Tang et al., 1996). Finally, the C-terminal region, which contains two plakin repeat domains, binds to keratin intermediate filaments (Fontao et al., 2003).

Recruitment of BP230 into HDs requires the association of plectin with α6β4 and the presence of BP180 (Koster et al., 2003), suggesting that BP230 is incorporated in the final stages of HD assembly (Wilhelmsen et al., 2006). Genetic mutations in the *DST* gene that target BP230 cause a mild form of epidermolysis bullosa simplex, a blistering disease characterized by skin fragility (Groves et al., 2010). Ablation of BP230 in mice resulted in a similar phenotype of blistering caused by mechanical stress (Guo et al., 1995). In the absence of BP230, HDs lack an intracellular substructure called the inner plaque, and the bundles of keratin filaments do not attach to the HDs. The role of BP230 in the attachment of intermediate filaments is further supported by the absence of the inner plaque in type II HDs and the less robust connection of intermediate filaments to type II than to type I HDs (Uematsu et al., 1994).

HDs are dynamic complexes that disassemble when epithelial cells migrate, for example during wound healing (Gipson et al., 1993) and in invasive carcinoma cells (Herold-Mende et al., 2001). In keratinocytes, epidermal growth factor and phorbol myristate acetate promote HD disassembly by inducing phosphorylation of the β4 subunit through the activation of the Ras/ERK-1/2 and protein kinase C (PKC) pathway (Frijns et al., 2010; Margadant et al., 2008; Rabinovitz et al., 1999). Phosphorylation of Ser residues in the CS (Frijns et al., 2010; Rabinovitz et al., 2004; Wilhelmsen et al., 2007) and in the C-tail (Frijns et al., 2012) of β4 disrupts the interaction with plectin and is a major mechanism for HD disassembly. Phosphorylation of β4 at S1424 correlates with a loss of co-localization with BP230 and BP180 (Germain et al., 2009); yet, little is known about the mechanisms that regulate the interaction of α6β4 with BP proteins during HD disassembly.

The structural understanding of protein-protein interactions in HDs is limited to the primary contact between α6β4 and plectin (de Pereda et al., 2009; Song et al., 2015). The isolated region FnIII-3,4 of β4 is reluctant to crystallize, thus, we had solved its structure using hybrid methods (Alonso-Garcia et al., 2015). Nonetheless, how the FnIII-3,4 engages with BP230 or other proteins remained unknown. Here, we present a detailed mapping of the mutual binding sites in β4 and BP230, the 3D structure of the β4-BP230 complex, and the conformational changes that binding causes in β4. Finally, we have identified potentially phosphorylatable residues in BP230 and β4 that play key roles in their interaction.

## Results

### Integrin β4 binds to a segment of the N-terminal tail of BP230

Residues 1-56 of BP230 interact with the FnIII-3,4 of β4 in yeast two-hybrid assays; and the longer segment 1-92 associated more efficiently with β4 (Koster et al., 2003). The region 1-92 includes the N-tail (residues 1-55) and the first a-helix of the SR2 (BP230 does not have the SR1). To investigate the binding to β4 *in vitro,* we created constructs of BP230 that include the complete SR2 (56-162) to maintain the integrity of the SR fold. The fragment 1-162 of BP230 had extremely low solubility that hampered its characterization. The longer construct of this region suitable for analysis was the 10-162. Using size exclusion chromatography (SEC), interaction was detected between β4 1436-1666 (β4-CS-FnIII-3,4) and BP230 10-162, but not with the isolated SR2 (Figure S1). Thus, direct binding of BP230 to β4 requires the N-tail.

In order to map the regions within the N-tail of BP230 responsible for binding to β4, the affinity of β4-CS-FnIII-3,4 for a series of BP230 N-terminal deletion mutants was determined (Figure 1B,D). β4 bound to BP230 constructs 10-162, 19-162, and 28-162 with similar affinity, suggesting that the segment 10-27 does not contribute to the interaction. The fragments of BP230 32-162 and 37-162 had slightly lower affinity with respect to the longer constructs. Affinity for BP230 41-162 was notably lower (5-fold larger equilibrium dissociation constant, *Kd*). Finally, BP230 46-162 and 56-162 did not bind to β4. In summary, residues 41-45 of BP230 are essential for binding to β4, and residues 28-40 also contributed to the interaction.

Despite the fact that the SR2 of BP230 was not sufficient to sustain binding to β4, we explored if it contributes to the interaction. First, we created a chimeric protein in which the SR2 of BP230, residues 59-162, was replaced by the SR2 of plectin, residues 420-530. β4-CS-FnIII-3,4 bound to the BP230/plectin chimera with a similar *Kd* as to the equivalent BP230 fragment 19-162, suggesting that the SR2 is not required for binding to β4 or that its contribution can be mimicked by the SR2 of plectin.

Next, we analyzed the binding of β4 to three synthetic peptides of the N-tail (Figure 1C,D). β4 bound to the BP230 peptide 26-55 with slightly higher affinity than to the BP230 fragment 28-162 that contains the SR2; further supporting the notion that the SR2 is dispensable for the interaction. Binding of β4 to two shorter BP230 peptides, 26-47 and 37-55, shows a 5-fold and a 3-fold increase in the *Kd* with respect to the 26-55 peptide, respectively. Thus, the segments 26-36 and 48-55 contribute to the interaction. Taking together, our data indicate that the region 26-55 of the N-tail of BP230, which is only present in this isoform of BPAG1, is sufficient for binding to β4.

### Mapping the BP230-binding site in β4

First, we analyzed the contribution to the binding to BP230 of the distinct modules included in the BP230-interaction region previously identified using yeast two-hybrid assays (Koster et al., 2003) (Figure 2A). The fragment β4-FnIII-3,4 (1457-1666) bound to BP230 with the same affinity as the longer β4-CS-FnIII-3,4, suggesting that the CS does not participate directly in the interaction. The individual FnIII domains did not bind to BP230, indicating that the FnIII-3,4 pair is necessary and sufficient for binding. In the yeast two-hybrid assays the CS of β4 might have been required for binding to BP230 to reduce steric hindrance between the Gal4 activation domain and the FnIII-3,4 of β4 in the fusion protein and to allow the correct transcriptional activation upon binding to BP230 fused to the Gal4 binding domain.

**Figure 2.**
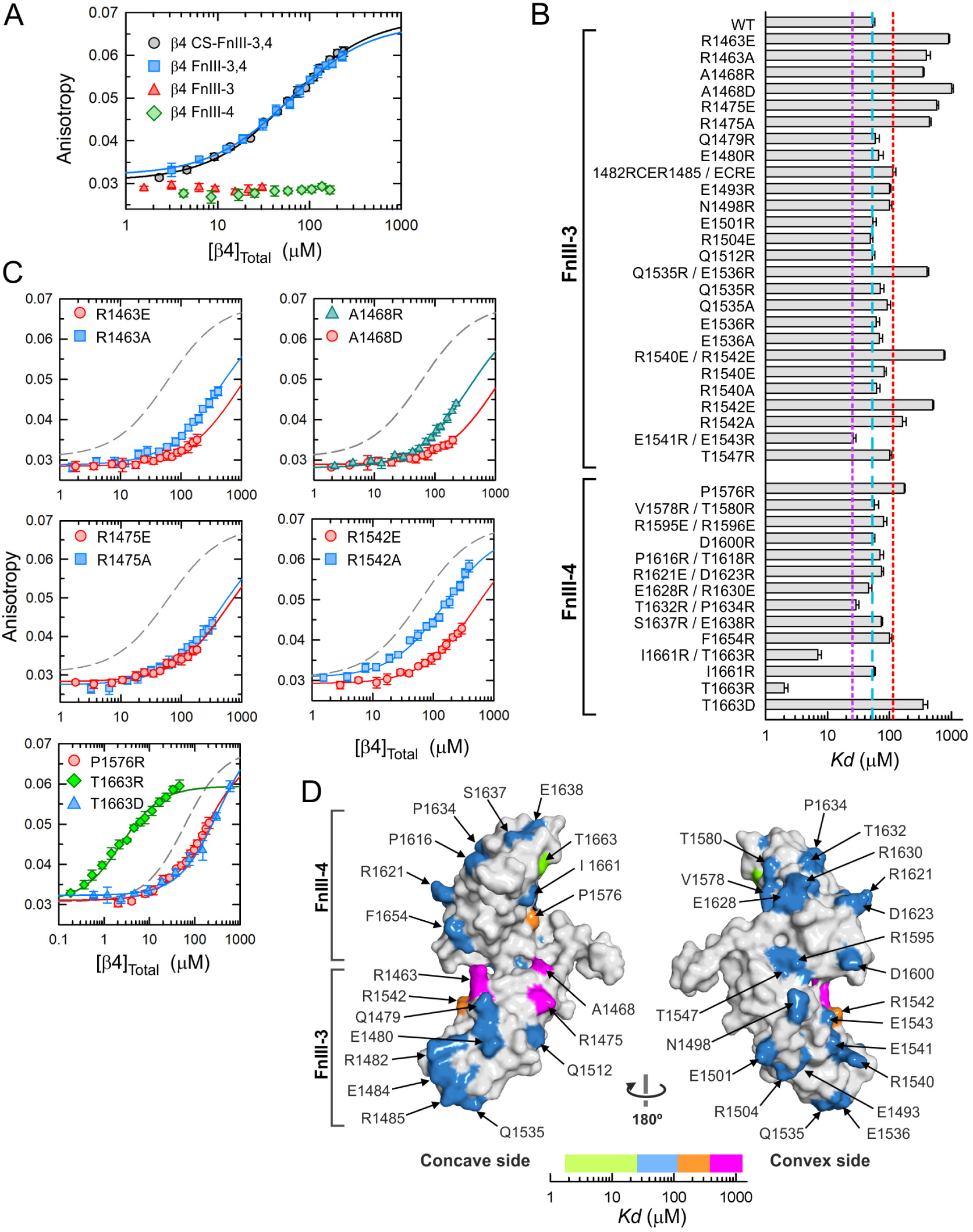
Mapping the BP230-binding site in β4. **(A)** Equilibrium binding of fragments of β4 to the fluorescein-labeled BP230 peptide 26-55 measured by fluorescence anisotropy. Lines represent the fit to the data. **(B)** Bar-chart of the *Kd* of the binding to BP230 26-55 of β4-CS-Fn-3,4, WT and mutants. Error bars show the asymptotic standard error. Dashed vertical lines mark the *Kd* of WT β4 (cyan) and the two-fold lower and higher values with respect to the WT (purple and red). **(C)** Equilibrium binding of representative mutants. The binding curve of WT β4 (dashed lines) is shown for comparison. **(D)** Surface representation of the unbound structure of β4-FnIII-3,4. Residues are colored according to the effect of their mutations on the affinity for BP230, as indicated by the color bar. For residues changed in double and single mutants, the effect of the latter is shown.

Next, we combined structure-based site directed mutagenesis with the fluorescent-based quantitative assay to map the BP230-binding site in β4 (Figure 2B,C). To prevent distortions of the FnIII fold, mutations were introduced at solvent-exposed residues in the structures of the FnIII-3 and FnIII-4 (Alonso-Garcia et al., 2015). Initially, we mainly created reverse-charge substitutions and, when possible, we changed two or three adjacent residues. The mutants Q1479R, E1480R, R1482E/E1484R/R1485E, E1493R, N1498R, E1501R, R1504E, Q1512R, E1541R/E1543R, and T1547R, which carry substitutions in the FnIII-3, showed values of the *Kd* within a 2-fold range, higher or lower, with respect to the wild type (WT) β4 protein, and were considered to have a small effect on the interaction. On the other hand, the mutants R1463E, A1468R, A1468D, R1475E, Q1535R/E1536R, and R1540E/R1542E showed a notable reduction in the affinity for BP230. The effects of reverse-charge substitutions could be caused by either the loss of contacts mediated by the WT side-chain, or by contributions of the engineered residue. To differentiate these two mechanisms, when a mutation altered significantly the affinity for BP230, we also analyzed the change to Ala. R1463A, and R1475A reduced the affinity for BP230, indicating that these two Arg play important roles for binding. To further characterize the double mutants that affect the binding, we also analyzed the individual substitutions. The single mutants Q1535R, Q1535A, E1536R, and E1536A showed a similar affinity for BP230 as the WT β4, suggesting that individually these two residues have a minor contribution to the binding. As for the pair R1540E/R1542E, the single mutant R1540E displayed only a minor effect on the binding, while R1542E significantly reduced the affinity for BP230. Similarly, R1542A, but not R1540A, reduced the affinity.

Similarly, we analyzed the effect of substitutions of residues in the FnIII-4. The β4 mutants V1578R/T1580R, R1595E/R1596E, D1600R, P1616R/T1618R, R1621E/D1623R, E1628R/R1630E, T1632R/P1634R, S1637R/E1638R, and F1654R had a similar affinity for BP230 as the WT protein. Only the substitution P1576R resulted in an increase in the *Kd* larger than 2-fold compared to the WT. The double mutant I1661R/T1663R had increased affinity for BP230 (∼8-fold reduction in the *Kd*). This effect was due to the change T1663R, while the mutant I1661R had a similar affinity for BP230 as the WT β4. In contrast to the effect of T1663R, the mutant T1663D showed reduced affinity for β4, further supporting a role of T1663 in the interaction.

We mapped the effect of the mutations onto the structure of the isolated FnIII-3,4 (Figure 2D). The two FnIII domains are arranged in a slightly bent manner forming a curved shape with concave and convex sides, and the inter-domain linker protrudes on one side (Alonso-Garcia et al., 2015). The mutations were widely distributed throughout the surface of the FnIII-3,4. Yet, the residues whose substitutions mostly altered binding to BP230 (R1463, A1468, R1475, R1542, P1576, and T1663) clustered at the centre of the concave side, suggesting that the BP230-binding area extends around the cleft formed by the two FnIII domains, which is evolutionary conserved (Alonso-Garcia et al., 2015).

### Structural characterization of the β4-BP230 complex

We used x-ray crystallography to unveil the structural basis of the BP230 binding to β4. The weak interaction between BP230 and β4 posed a major challenge to crystallize their complex. To overcome this limitation, the structure of the β4-BP230 complex was initially elucidated using the high-affinity mutant β4-T1663R bound to BP230 26-55; this structure was solved to 1.55 Å. Subsequently, the structure of the β4(WT)-BP230 complex was solved to 2.05 Å. The WT and mutant complexes crystallized in the same crystal form that contains a single copy of the complex in the asymmetric unit (Table 1 and Figure 3). The two structures were almost identical; after superimposition, the root mean square displacement (rmsd) of all main-chain atoms between the two structures was 1.03 Å. Only a segment of the inter-domain linker of β4, residues 1557-1564, shows differences between the two structures (see below). Hereafter, the WT structure is described unless otherwise indicated.

**Table 1.** Crystallographic data collection and refinement statistics

| Complex | $\beta 4$(T1663R)-BP230 | $\beta 4$(WT)-BP230 |
| --- | --- | --- |
| **Data Collection** |  |  |
| X-ray beamline | Xaloc (Alba) | i03 (Diamond) |
| Space group | C2 | C2 |
| Cell dimensions | a = 105.6 Å<br>b = 59.5 Å<br>c = 42.4 Å<br>$\beta$ = 113.5° | a = 104.5 Å<br>b = 60.7 Å<br>c = 40.8 Å<br>$\beta$ = 113.7° |
| Wavelength (Å) | 0.97915 | 0.97625 |
| Resolution (Å) | 1.55 (1.59-1.55)<sup>a</sup> | 2.05 (2.10-2.05)<sup>a</sup> |
| Unique reflections | 35010 (2562)<sup>a</sup> | 14766 (1091)<sup>a</sup> |
| Average multiplicity | 19.9 (19.4)<sup>a</sup> | 6.7 (6.9)<sup>a</sup> |
| Completeness (%) | 99.7 (99.8)<sup>a</sup> | 99.5 (100)<sup>a</sup> |
| R<sub>meas</sub><sup>b</sup> (%) | 5.6 (350)<sup>a</sup> | 10.5 (195)<sup>a</sup> |
| CC 1/2 (%) | 100 (70.5)<sup>a</sup> | 99.8 (69.4)<sup>a</sup> |
| Mean I/σI | 26.4 (1.53)<sup>a</sup> | 11.7 (1.28)<sup>a</sup> |
| **Refinement** |  |  |
| Resolution range (Å) | 39 – 1.55 | 48 – 2.05 |
| Unique reflections, work/free | 33244 / 1749 | 14019 / 703 |
| R work (%) | 19.5 | 22.4 |
| R free<sup>c</sup> (%) | 21.0 | 23.6 |
| Number of |  |  |
| residues ($\beta 4$ / BP230) | 202 / 24 | 199 / 23 |
| waters | 105 | 46 |
| Glycerol | 2 | - |
| Average B value (Å<sup>2</sup>) |  |  |
| Wilson plot | 31.6 | 45.8 |
| Protein ($\beta 4$ / BP230) | 44.4 / 42.5 | 64.3 / 63.2 |
| Solvent | 41.7 | 48.7 |
| Glycerol | 51.3 | - |
| rmsd bond lengths (Å) | 0.004 | 0.002 |
| rmsd angles (°) | 0.721 | 0.496 |
| Ramachandran distribution<sup>d</sup> |  |  |
| Favored regions | 212 (97.7%) | 213 (98.2%) |
| Additionally allowed | 5 (2.3%) | 4 (1.8%) |
| Outliers | 0 | 0 |
| PDB code | 6GVK | 6GVL |

<sup>a</sup> Numbers in parenthesis correspond to the outer resolution shell.<sup>b</sup> R<sub>meas</sub> is the multiplicity independent R factor as described (Diederichs and Karplus, 1997).<sup>c</sup> Calculated using 5% of reflections that were not included in the refinement.<sup>d</sup> Analyzed with the program MOLPROBITY (Davis et al., 2007).

**Figure 3.**
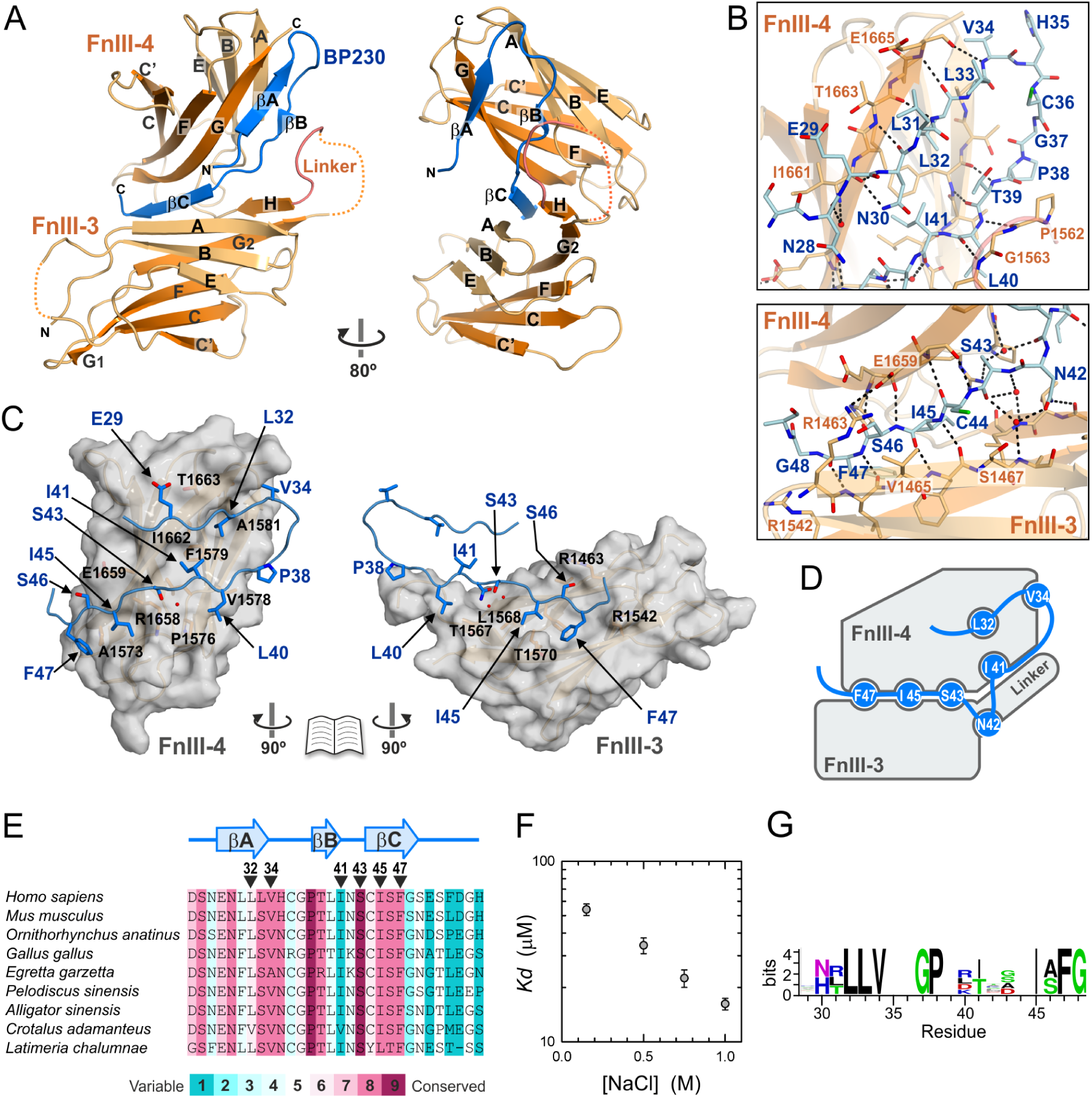
Structure of BP230 bound to the FnIII-3,4 of β4. **(A)** Two views of a ribbon representation of the structure of the WT β4-BP230 complex. **(B)** Close up of contacts between BP230 (blue) and β4 (orange); waters are shown as red spheres. Polar contacts are shown as dashed lines. **(C)** Open-book view of the footprint of BP230 on the surface of the FnIII-4 (left) and the FnIII-3 and the liker (right) of β4. **(D)** Schematic illustration of the BP230 residues that dock into pockets of β4. **(E)** Alignment of the region 26-55 of human BP230 with the sequences of representative species, colored by the evolutionary conservation. A multiple alignment of the sequences of 36 species used to calculate the conservation scores is shown in the supplementary Fig S6. **(F)** Dependence of the *Kd* of the BP230-β4 interaction on the salt concentration.

The structures of the FnIII-3 and FnIII-4 in the complex are very similar to those observed in the crystals of the isolated domains (Figure S2). The inter-domain linker of β4, which was only approximately modelled in the structure of the free FnIII-3,4, contributes to the organization of the two FnIII domains and to the BP230-binding interface. The final part of the linker, residues 1566-1569, form a β-strand H that lies between the two FnIII domains and participates in the BP230-binding site. Residues 1562-1564, upstream strand H, contact BP230 making two main-chain H-bonds. In the β4(T1663R)-BP230 complex, the linker also contacts BP230, but it adopts a slightly different conformation (Figure S3). T1663R is unlikely to cause these differences in the linker because it does not induce local changes in BP230.

Continuous electron density was observed for residues 27-49 of BP230 in the WT structure and for residues 27-50 in the T1663R structure (Figure S4). BP230 adopts an extended hairpin-like structure that contains three β-strands (βA, βB, and βC) (Figure 3A). The first part of BP230 (residues 27-41) packs against one side of the FnIII-4; strands βA (29-33) and βB (39-40) make β-contacts with strands G2 and A of the FnIII-4, respectively (Figure 3B). βC (44-48) makes simultaneous backbone H-bonds with the strands A of the FnIII-3 and G2 of the FnIII-4, creating a continuous β-sheet that extends along the two FnIII domains. The β-strands of BP230 in the complex are stabilized by contacts with β4. Yet, in the absence of β4 the N-tail is highly sensitivity to controlled proteolysis (Figure S5), suggesting that this region of BP230 is mostly unstructured on its own.

The side chains of several residues of BP230 also contribute to the binding interface (Figure 3B-D). L32 and I41 are buried in hydrophobic pockets on the FnIII-4. S43, I45, and F47 dock in pockets formed at the interface of the two FnIII domains. V34 and L40 sit on shallower cavities on the surface of β4. Most of the residues that bind in the pockets of β4 are highly conserved in multiple species (Figures 3E and S6). In addition, the side chains of N30 and N42 make H-bonds to the carbonyls of I1661 and F1566, respectively. In summary, binding is mainly driven by hydrophobic contacts, in accordance with an increase in the affinity of the interaction at high salt concentrations (Figure 3F).

To gain further insight of the determinants of the BP230-recognition by β4, we applied a structure-base computational method to predict the BP230 sequences tolerated at the interface (Figures 3G and S7). In addition to the residues that dock into the β4 pockets, high preference is predicted for G37 and P38, which is in agreement with their conservation. Of these two residues, only P38 contacts β4. Glycines appear frequently before prolines, where they facilitate the cis-trans isomerization of the latter, suggesting that the G37-P38 tandem might play an important role in the local rearrangement of the BP230 during binding to β4. Noteworthy, BP180 and Solo, which interact with the FnIII-3,4 region of β4, do not have sequence similarity with the tolerated binding motif.

The crystal structure is in good agreement with the mutagenesis data (Figure S8). R1463 in the FnIII-3 forms a salt bridge with E1659 in the FnIII-4 that closes over BP230. The inhibitory effect of the R1463E and R1463A mutations suggests that this inter-domain clasp is essential to stabilize the interaction. A1468 is buried facing the β4 inter-domain linker; an Arg or Asp at this position may clash with the linker altering the binding site. R1475, located near the inter-domain cleft that accommodates the strand βC of BP230, does not engage directly with BP230, suggesting that the substitutions R1475E and R1475A affect indirectly the binding site. R1542 is near G48 of BP230; in addition, its side chain makes an H-bond with P1461 and contributes to the positioning of T1462, which in turn contacts BP230. Hence, the R1542E and R1542A substitutions are likely to alter this region of the interface. On the other hand, the nearby R1540 does not contact BP230, which explains the lack of effect on the binding of the R1540E and R1540A changes. P1576 in the FnIII-4 contacts L40 of BP230; the P1576R substitution is likely to reduce binding by distorting the interface. Finally, T1663 is near E29 of BP230. In the structure of the β4(T1663R)-BP230 complex this engineered Arg makes an additional salt bridge with E29 without altering the conformation of E29 or the rest of BP230, which explains the increased affinity for BP230 of the T1663R mutant. On the other hand, the T1663D substitution is likely to reduce the affinity by creating an electrostatic repulsion with BP230.

Since mutations in β4 that affect binding to BP230 are near the inter-domain interface, we analyzed their effect on the structure of the FnIII-3,4 using small angle x-ray scattering (SAXS) (Figure S9 and Table S1). Mutants R1463E, R1463A, R1475A, R1542E, R1542A, and T1663R behaved as the WT protein. The mutant R1475E showed slight differences with the WT protein, suggesting moderate self association. Only mutations A1468R and A1468D induced large deviations with respect to the WT β4 fragment, which correspond to a large inter-domain flexibility. A1468 does contact BP230 directly. Thus, the deleterious effect of A1468R and A1468D on binding might be caused by the disruption of the inter-domain arrangement, suggesting that BP230 recognizes a pre-ordered surface in β4.

### Binding of BP230 induces a conformational change in β4

Superimposition of the FnIII-3 domains in the structures of FnIII-3,4 bound to BP230 and in the free form, revealed differences in the relative orientation of the FnIII-4 domain (Figure 4A). The orientations of the FnIII-4 in the two states are related by a ∼38° rotation around an axis located along the inter-domain interface.

**Figure 4.**
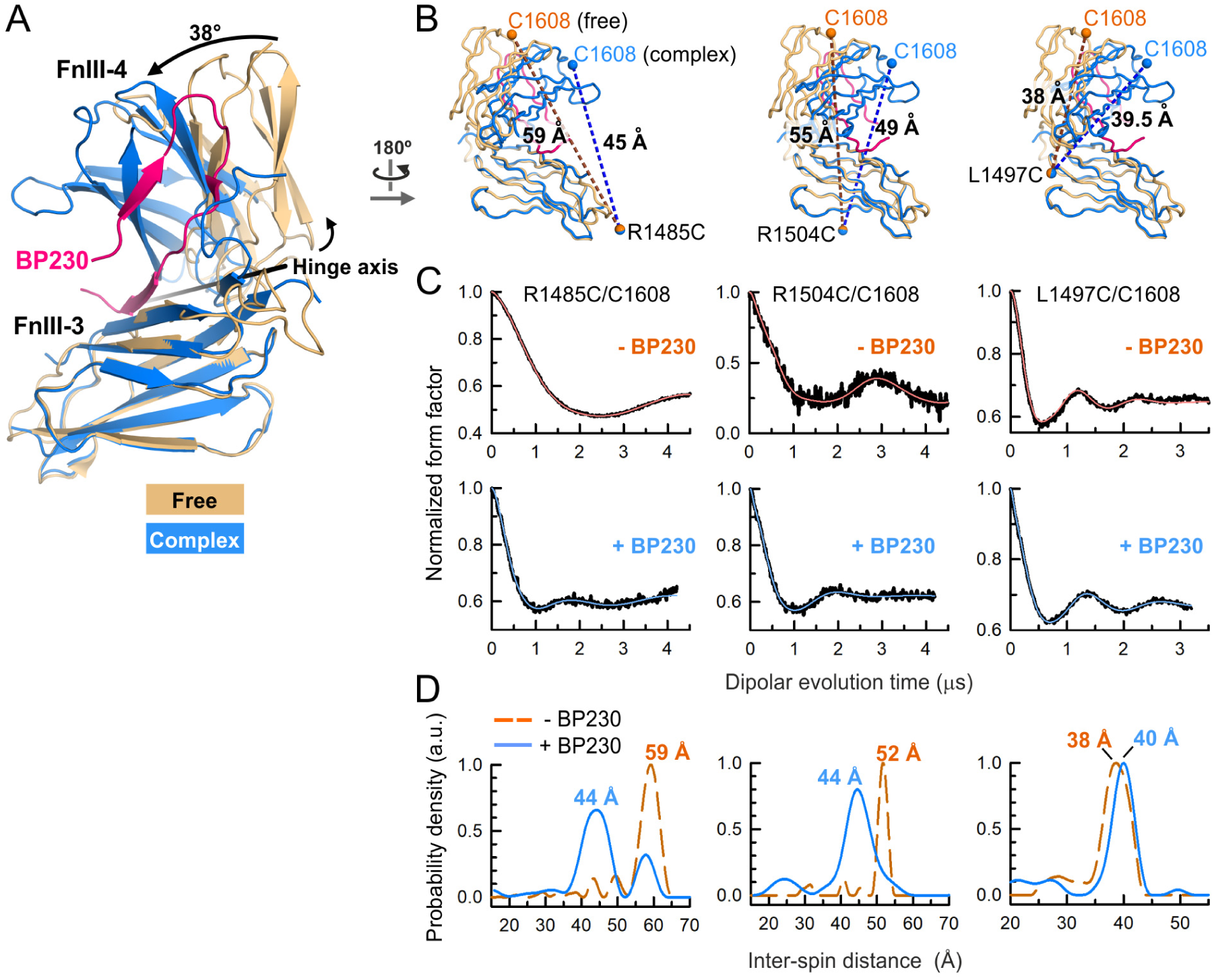
BP230 induces a conformational change in the FnIII-3,4 of β4. **(A)** Comparison of the structures of β4 FnIII-3,4 in the free form (orange) and bound to BP230 (β4 blue and BP230 magenta). Only the Cα atoms of the FnIII-3 were used for the superimposition. The orientations of the FnIII-4 in the two structures are related by a rotation around a hinge axis (black line). **(B)** Worm models of the two structures in A. The modeled average positions of the MTSL-paramagnetic centers (spheres) attached to the Cys pairs and the distances (dashed lines) in the two conformations are shown. **(C)** DEER normalized dipolar evolution (black lines) and fits to the data for the MTSL-labeled β4 mutants in the absence (top) and in the presence of BP230 26-55 (bottom). **(D)** Inter-spin distance distributions calculated from the data in C.

To characterize the conformational change in β4 induced by BP230, we used Double Electron-Electron Resonance (DEER) spectroscopy, a method that was useful to measure long range inter-domain distances between engineered paramagnetic groups in the FnIII-3,4 (Alonso-Garcia et al., 2015). We used β4-FnIII-3,4 mutants that contain the WT C1608 in the FnIII-4 and a second Cys introduced in the FnIII-3 at R1485C, R1504C, or L1497C (Figure 4B). Proteins were doubly labeled with the spin-probe MTSL and the distances between the pairs of paramagnetic groups were measured in the absence and in the presence of BP230 26-55 (Figure 4C,D). The distances in the free state matched those previously observed (Alonso-Garcia et al., 2015). In the presence of BP230, the peaks of the inter-spin distance distributions changed and were in agreement with the distances modeled in the β4-BP230 complex. In summary, BP230 induces closing of the FnIII domains onto BP230, with the inter-domain linker acting as a hinge.

### The β4-BP230 interaction is required for the recruitment of BP230 into HD

To assess the role of the binding interface on the recruitment of BP230 into HDs, we analyzed the distribution of BP230 and other hemidesmosomal proteins in PA-JEB keratinocytes (that do not express endogenous β4) in which WT or point mutants of β4 were expressed (Figure 5). Endogenous BP230 co-localized with WT β4 and with endogenous BP180 and plectin, displaying a punctuated pattern characteristic of HDs. In contrast, BP230 had a diffused localization in PA-JEB keratinocytes expressing β4 R1463E or R1463A, and BP230 did not co-localize either with BP180 or with plectin. β4, BP180, and plectin showed a patched distribution in PA-JEB/β4-R1463E or R1463A cells, which supports the view that BP230 is not required for the incorporation of these proteins into HDs (Koster et al., 2003; Wilhelmsen et al., 2006). When the high-affinity β4 mutant T1663R was expressed, BP230 was again recruited into HDs and colocalized with β4, BP180, and plectin. In summary, the interaction between the N-tail of BP230 and the FnIII-3,4 is a major determinant for the incorporation of BP230 into HDs in keratinocytes.

**Figure 5.**
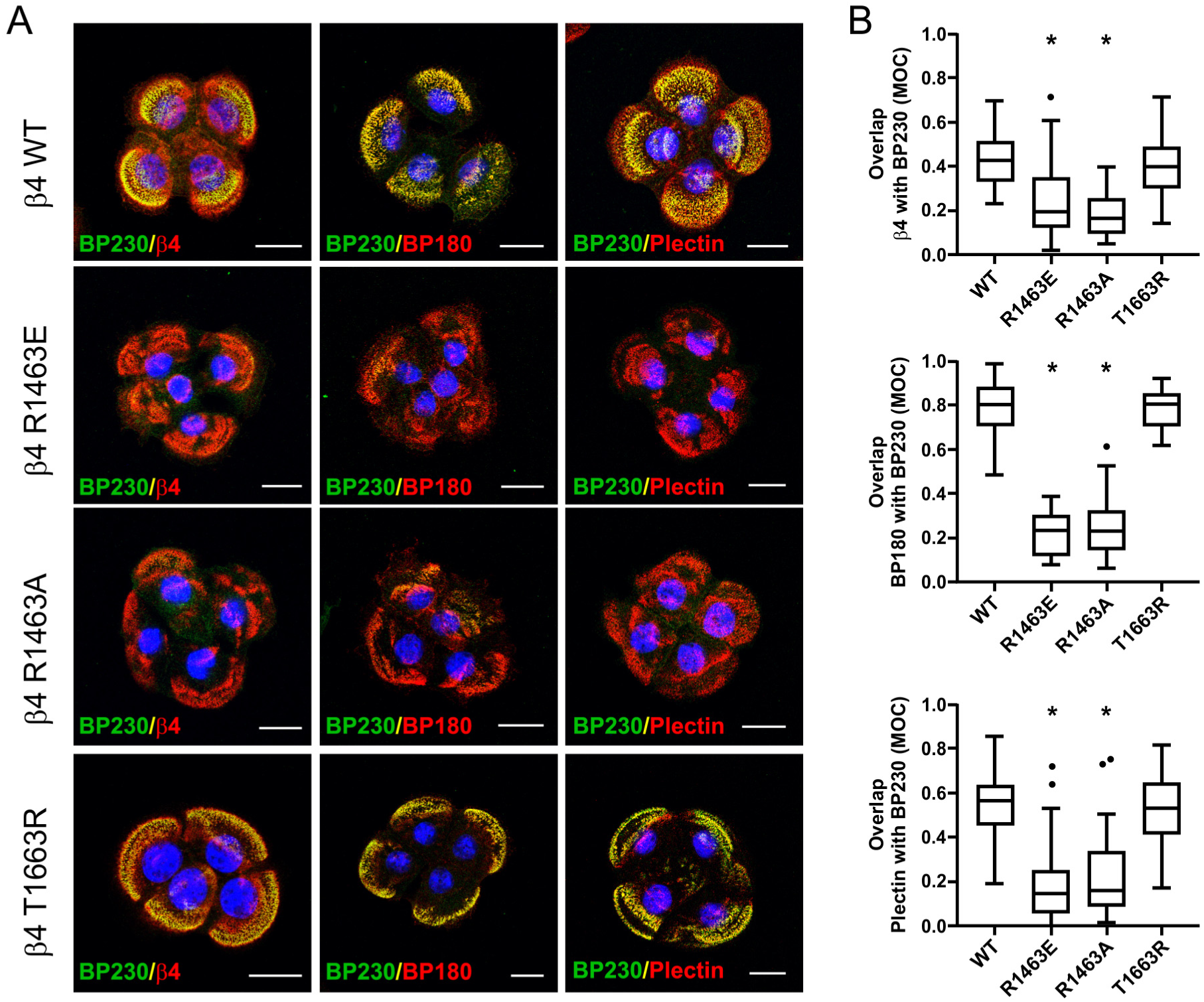
Binding of BP230 N-tail to β4 FnIII-3,4 is necessary for the recruitment of BP230 into HDs. **(A)** Confocal microscopy images of PA-JEB keratinocytes stably expressing β4 WT or the mutants R1463E, R1463A, or T1663R. Cells were stained with antibodies against BP230 (green), β4, BP180, or plectin (red), as indicated. Co-localization of BP230 with the other HD proteins in the presence of β4 WT or the high-affinity mutant T1663R appears as yellow. BP230 is diffusely distributed in the presence of the low affinity β4 mutants R1463E and R1463A. Nuclei were stained with DAPI (blue). Scale bars: 20 μm. **(B)** Quantification of the co-localization of β4, plectin or BP180 with BP230. Manders’ overlap coefficients (MOC) were calculated for at least 14 images per condition (14-72 images per condition from 1-3 experiments). Tukey box plots show the median (solid line), the 25^th^ and 75^th^ percentiles (boxes) for each distribution; whiskers represent 1.5 times above or below the interquartile range, outlier points outside the whiskers are displayed (dots). The P-values (* = P < 0.0001) were calculated using the unpaired, nonparametric Mann-Whitney test.

### Role of potentially phosphorylatable residues in the interaction

Phosphorylation of β4 by Ser/Thr kinases promotes HD disassembly by inhibiting the interaction of the integrin with plectin. This prompted us to analyze the role of potentially phosphorylatable residues in the binding of BP230 to β4. The region FnIII-3,4 of β4 contains 21 predicted putative phosphorylatable Ser/Thr residues. Of those, only T1663 is in direct contact with BP230 (Figure S10). As described above, the phosphomimetic change T1663D reduced the affinity for BP230 (Figure 2B,C). Another predicted phosphorylatable residue of β4, S1556, is located in the inter-domain linker and it was disordered in the crystal structure. The sequence context of S1556 (PQSP) fits the optimal substrate sequence of the ERK1/2 kinases (P-X-S/T-P), which are known to phosphorylate S1356 in the CS leading to the dissociation from plectin (Frijns et al., 2010). Yet, the substitution S1556D did not affect the affinity for BP230 with respect to WT β4 (Figure S10B).

We also analyzed the role of potentially phosphorylatable residues in BP230 on the interaction with β4 (Figure 6A-C). BP230 has six Ser/Thr residues in or near the β4-binding site. The phosphomimetic substitutions S27D, S43D, S49D, and S51D did not affect the affinity for β4. The substitution T39D resulted in a ∼3-fold increase in the *Kd.* Finally, the BP230 mutant S46D showed the weakest binding to β4. Analysis of the effect of these point mutations on the interaction with β4 using a pull-down assay (Figure 6C) showed that the BP230 mutants S27D, S43D, S49D, and S51D bound to GST-β4-CS-FnIII-3,4 similarly as the WT protein. Only a faint amount of the mutant T39D bound to β4, and no interaction was observed for the mutant S46D. Thus, there is a good agreement between the results obtained with these two orthogonal methods.

**Figure 6.**
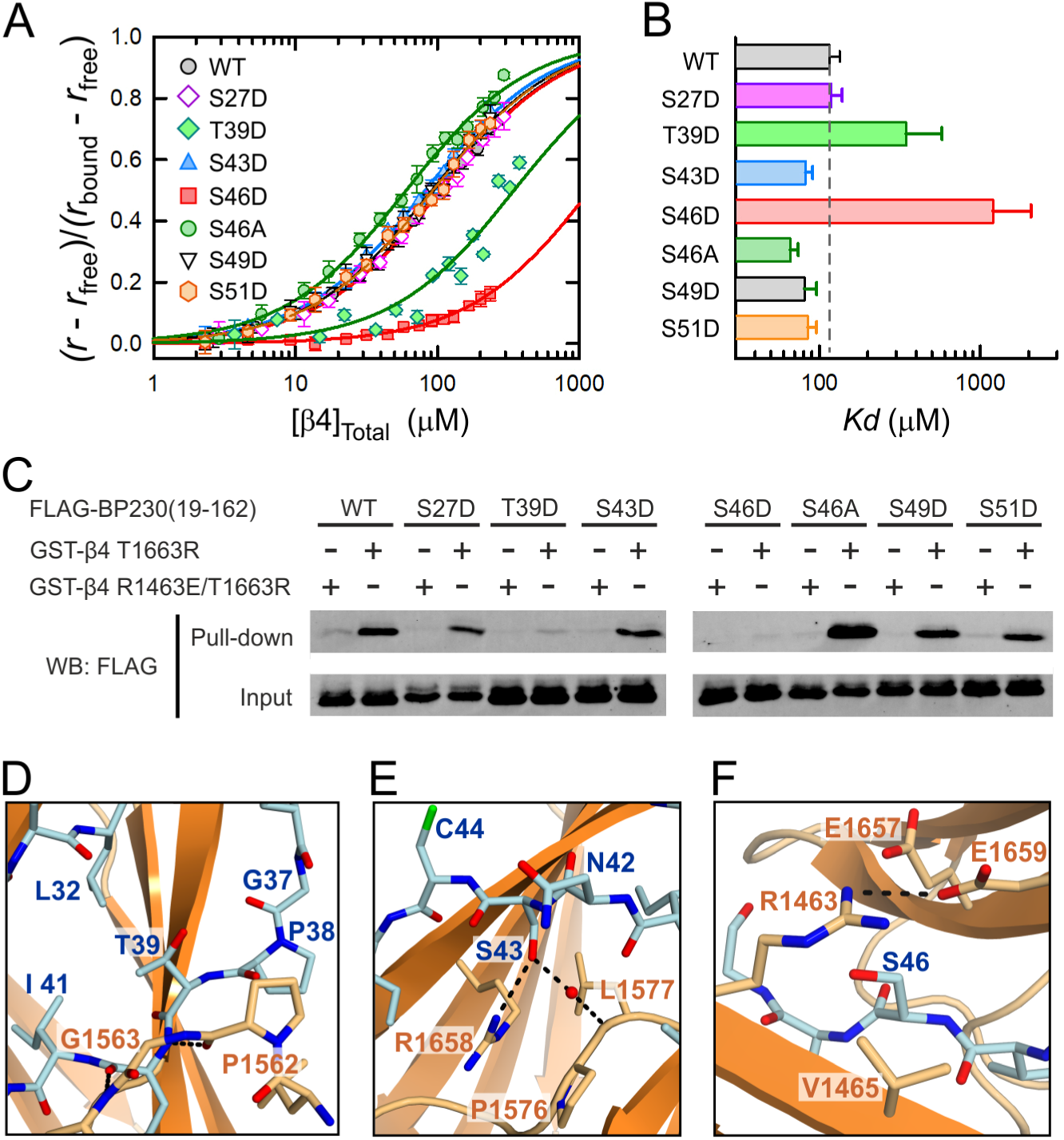
Effect of phosphomimetic mutations in BP230 on the binding to β4. **(A)** Equilibrium binding of β4-CS-FnIII-3,4 to the BP230 10-162 fragment, WT and point mutants, measured by fluorescence anisotropy. Lines represent the fit to the data. **(B)** Bar-chart of the *Kd* values in A. Error bars show the SD (2-3 experiments). **(C)** Pull-down analysis of the interaction between FLAG-BP230 (19-162), WT and point mutants, and GST-β4 (1436-1666)-T1663R. The BP230 binding-deficient double mutant R1463E/T1663R was used as a negative control. BP230 proteins in the cell lysates (input) and in the bound samples (pull-down) were analyzed by WB with an anti-FLAG antibody. **(D-F)** Detail of the structural environment of T39 (D), S43 (E) and S46 (F) of BP230 in the complex.

S27, S49 and S51 are at the ends of the β4-binding sequence. T39 is adjacent to the inter-domain linker of β4 (Figure 6D); hence, the substitution T39D could disturb contacts with β4. S43 docks in a polar pocket of β4 that can accommodate an acidic group (Figure 6E). S46 is covered by the inter-domain salt bridge formed by R1463 and E1659 of β4 (Figure 6F). The S46D substitution is likely to inhibit binding to β4 by interfering with the R1463-E1659 salt bridge, which is in agreement with the inhibitory effect of the R1463A substitution. The mutant S46A showed slightly higher affinity for β4 (Figure 6A-C); supporting the idea that the hydroxyl of S46 is not required for binding and that it imposes a penalty on the interaction. Collectively, our findings point to a potential role of T39 and S46 of BP230 and T1663 of β4 in the regulation of the interaction.

## Discussion

Integrin α6β4 binds to plectin and BP230 through similar regions, formed by pairs of tandem FnIII domains, but these interactions occur through very distinct mechanisms. The FnIII-1,2 region of β4 binds to a fairly flat surface on the ABD of plectin, which is a globular domain. In this study, we show that the FnIII-3,4 recognizes a linear motif in the N-tail of BP230, which has a relaxed structure in the free state. Our results suggest a mechanism for the high specificity and relatively low affinity of the β4-BP230 complex. The extensive binding interface that includes specific contacts provides the basis for a selective interaction. On the other hand, binding is likely to cause a disorder-to-order transition of the N-tail of BP230, which would have an unfavorable contribution that would moderate the affinity (Dyson and Wright, 2005). The low affinity of the β4-BP230 interaction is similar to that of the β4-plectin complex (*Kd* ∼30 μM). This supports the notion that HDs function as macromolecular Velcro^®^ fasteners, in which firm attachment is achieved by combining multiple weak contacts in a process mediated by the presence of multiple binding sites within each protein, and the homo-association and clustering of HD proteins.

Incorporation of BP230 into HDs requires the previous interaction between plectin and α6β4, and the recruitment of BP180. This is depicted in a hierarchical model for the assembly of HDs (Koster et al., 2003; Wilhelmsen et al., 2006). The mechanisms for the recruitment of BP230 to HDs only at the final stage of assembly remained largely uncharacterized. BP230 interacts with the cytoplasmic domains of β4 and BP180. The region 293-549 of BP230, which corresponds to the SR3-SR4-SH3-SR5 segment of its plakin domain, interacts with BP180, and this region of BP230 is required for the efficient recruitment into HDs (Koster et al., 2003). We have shown that point mutations in the FnIII-3,4 of β4 that disrupt the BP230-binding interface and the direct interaction *in vitro,* also prevent the recruitment of BP230 to HDs in keratinocytes in culture. This suggests that binding to β4, as observed in the 3D structure of the β4-BP230 complex, is essential for the incorporation of BP230 in the HDs.

Besides providing additional binding sites, BP180 or plectin might favor the β4-dependent recruitment of BP230 by regulating the affinity of the BP230-β4 interaction (Figure 7). The BP230-binding site is accessible in the isolated FnIII-3,4 fragment, and binding of BP230 causes a closure of the FnIII-3 and FnIII-4 domains. In the full-length α6β4, the cytodomain of β4 adopts a folded-over conformation in which the C-tail is positioned very close to the CS (Frijns et al., 2012), possibly through a direct intramolecular interaction between the CS and the C-tail (Koster et al., 2004; Rezniczek et al., 1998; Schaapveld et al., 1998). Given that the CS and C-tail flank the FnIII-3,4 region, this intramolecular clasp might mask the BP230-binding site in the FnIII-3,4 or might reduce the interdomain flexibility of β4 impeding the conformational change linked to the binding of BP230. Plectin binds to β4 at two sites. The ABD binds to the FnIII-1,2 and the beginning of the CS of β4, and this interaction induces a reorganization of that part of the CS (de Pereda et al., 2009). In a second site of contact, the plakin domain of plectin binds to the CS and the C-tail of β4. Thus, plectin binding to β4 might alter the organization of the CS and C-tail, and as a consequence that of the FnIII-3,4. BP180 might also affect the FnIII-3,4 region through its interaction with the FnIII-3. In summary, plectin and BP180 could also favor allosterically the interaction between BP230 and β4 by exposing the BP230-binding site or by reducing its rigidity.

**Figure 7.**
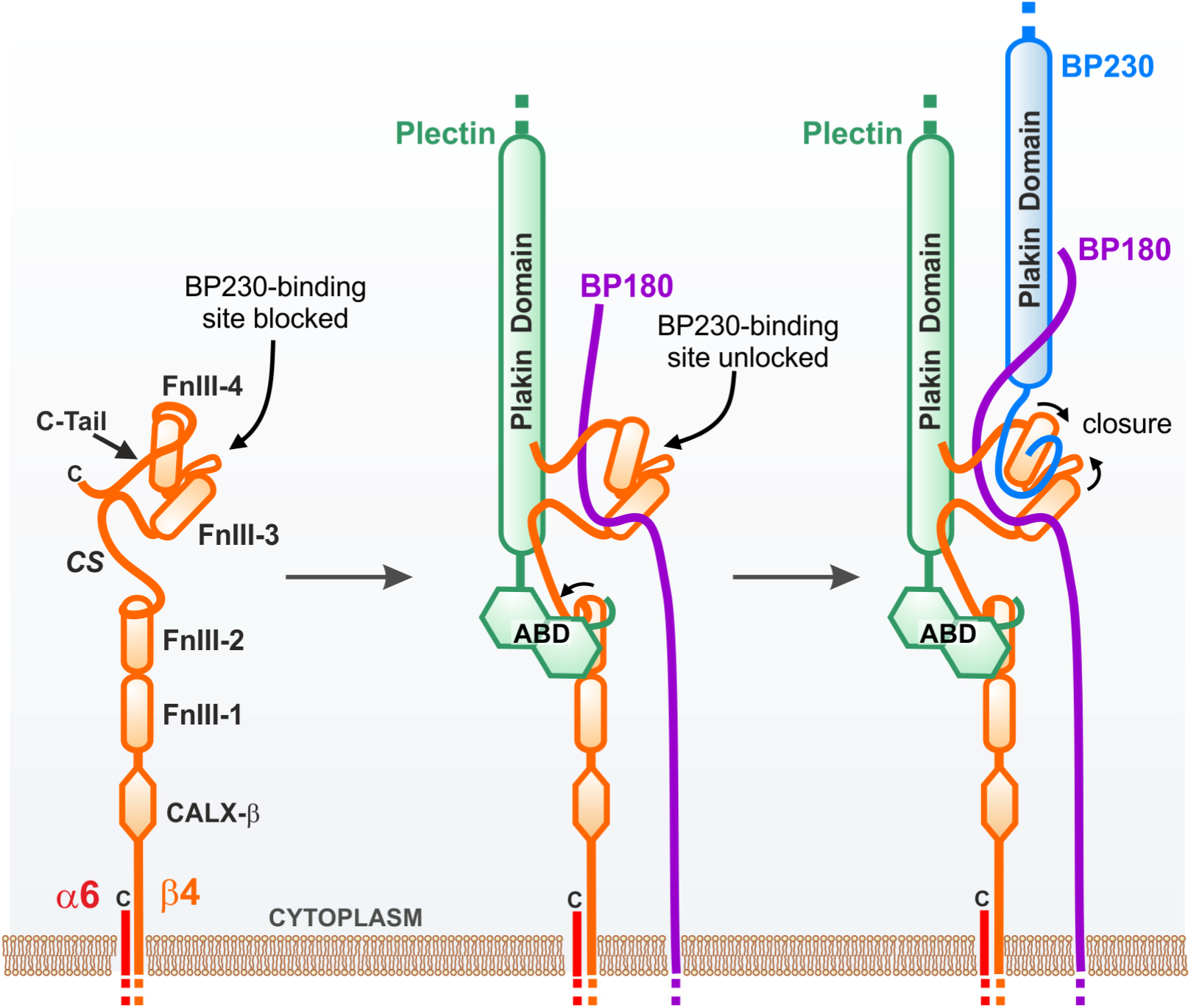
Model of regulation of BP230-β4 interaction during HD assembly. Schematic representation of three steps during HD assembly. Recruitment of BP230 might require unlocking of the BP230-binding site in β4, induced by plectin and/or BP180 binding to β4, and a second interaction between BP230 and BP180.

The regulation of the interaction between β4 and plectin, mainly through the phosphorylation of β4 at specific Ser and Thr residues, plays a major role in the disassembly of HDs in keratinocytes. Phosphorylation of S1356 and S1364 in the CS prevents the association with the ABD of plectin, and phosphorylation of T1736 in the C-tail of β4 reduces the interaction with the plakin domain of plectin (Frijns et al., 2012). It is unclear if other interactions in the HDs are directly inhibited during the disassembly, or if they are lost as a consequence of the disruption of the β4-plectin complex. Phosphorylation of S1424 in the CS of β4 is linked to a reduced association of β4 with BP180 in migrating keratinocytes (Germain et al., 2009). S1424 is located outside the region of β4 that interacts with BP180 (i.e. the FnIII-3) and it is not known if phosphorylation of S1424 reduces the affinity for BP180 directly. BP180 is also phosphorylated, probably at Ser/Thr, upon activation of PKC, which is linked to the dissociation of BP180 from HDs (Kitajima et al., 1999). Yet, it is not clear if the PKC-mediated phosphorylation of BP180 disrupts its interactions with other hemidesmosomal proteins and it is required for the mobilization of BP180 from HDs, or if PKC mobilizes indirectly BP180 through the phosphorylation of β4 and the disruption of the β4-plectin interaction.

We have identified three potentially phosphorylatable residues, T1663 in β4 and T39 and S46 in BP230, which are critical for the BP230-β4 interaction. Substitution of these residues with Asp severely reduced the interaction. Of those, the strongest inhibition was observed for the S46D mutant of BP230. S46 is highly conserved and it is only replaced by Thr in coelacanths (Figure 3E). The conservation of S46 is striking because the hydroxyl group destabilizes, albeit moderately, the binding to β4. This suggests that there has been a pressure to retain a Ser/Thr in this position. On the one hand, S46 could limit the affinity for β4, which in turn could allow for an indirect regulation of the BP230-β4 interaction during phases of HD assembly and disassembly. Alternatively, S46 might be a site for the direct regulation of the interaction. Phosphorylation of the N-tail of BP230 has not been described during HD disassembly; but its relaxed structure suggests that it is a preferential region for post-translational modification. Nonetheless, since several kinases act on β4 and BP180, it is reasonable that other proteins present in the HD niche might be phosphorylated as well. Future studies will be required to assess whether BP230 is post-translationally modified during the mobilization of hemidesmosomal proteins and whether active disruption of the BP230-β4 interaction plays a role in the disassembly of HD.

Finally, our structural and mechanistic description of the β4-BP230 interaction provides the basis to investigate its potential alteration in blistering diseases, and its regulation in normal keratinocytes during resting state, proliferation, and migration, and in invasive carcinoma cells.

## Materials and methods

### Protein expression and purification

The cDNA sequences coding for the fragments 1-162, 10-162, 19-162, 28-162, 32-162, 37-162, 41-162, 46-162, and 56-162 of human BP230 (Uniprot Q03001-8), and for the fragment 1436-1666 of human integrin β4 (Uniprot P16144-2) were cloned into a modified pET15b vector (pETEV15b) (Alonso-García et al., 2009). Plasmids encoding for the β4 fragments 1457-1666, 1457-1548, and 1572-1666 in pETEV15b were described earlier (Alonso-Garcia et al., 2015). The cDNA coding for a chimeric protein consisting of BP230 residues 19-58 and the SR2 of plectin, residues 420-530 (UniprotKB Q15149-2 was constructed by overlap extension PCR and was cloned in the pETEV15b vector as above. The cDNA of β4 (1436-1666) was also subcloned into a pGEX-4T3 vector that was modified to have restriction sites compatible with pET15b. Point mutations were introduced by PCR using the QuikChange method. The correctness of all constructs was verified by DNA sequencing.

Proteins were expressed in *Escherichia coli* strain BL21(DE3). The SR2 (56-162) of BP230 and the β4 proteins were produced soluble and were purified by affinity chromatography as described (Manso et al., 2016). Other BP230 recombinant proteins were produced insoluble; they were purified under denaturing conditions and were refolded by rapid dilution as described (Alonso-García et al., 2009). The His-tag was cleaved by digestion with TEV (tobacco etch virus) protease, unless otherwise indicated. GST-β4 (1436-1666) proteins were purified by affinity chromatography using a glutathione-agarose column.

### Peptides

Peptides corresponding to the regions of BP230 26-46, 37-55, and 26-55 were custom synthesized labeled with N-terminal fluorescein (Thermo Fisher Scientific). Unlabeled peptide 26-55 was obtained from Genosphere Biotechnologies.

### Labeling of proteins with Oregon Green 488

BP230 proteins were labeled in thiol groups with Oregon Green 488 iodoacetamide (Invitrogen). Proteins at 250 μM in 20 mM Tris (pH 8.0), 150 mM NaCl were incubated overnight at 4 °C with a 10-fold molar excess of the fluorescence probe. The reaction was stopped by adding a 10-fold molar excess of dithiothreitol (DTT) over the reagent and the proteins were separated from the unreacted probe by SEC using a Sephadex G25 (1 x 30 cm) column.

### Fluorescence-based binding assay

Labeled BP230 proteins and peptides at 1 μM in 20 mM Na-phosphate (pH 7.5), 150 mM NaCl, 1 mg/ml bovine serum albumin (BSA), unless otherwise indicated, were titrated with β4-CS-FnIII-3,4. The fluorescence anisotropy of the probe was measured with a FluoroMax-3 spectrofluorometer (HORIBA-Jobin-Yvon) equipped with Glan-Thompson polarizers, using 496 nm excitation wavelength and collecting the emission at 518 nm. The *Kd* and the fluorescence anisotropy of the free (*r*_*F*_) and fully bound (*r*_*B*_) states were estimated by fitting the following equation of a 1:1 binding model:

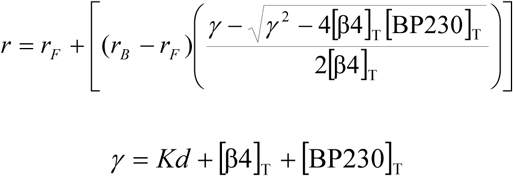

Where, [BP230]_T_ is the total concentration of labeled-BP230, [β4]_T_ is the total concentration of β4 added, and *r* is the observed fluorescence anisotropy of the probe.

### Crystallization and determination of the structure of the β4-BP230 complex

Samples of the β4-FnIII-3,4 (1457-1666) mutant T1663R in complex with the peptide of BP230 (26-55) were prepared by adding 300 μM of peptide from a 6 mM stock in dimethyl sulfoxide (DMsO) to a 210 μM (5 mg/ml) solution of β4-FnIII-3,4 in 10 mM Tris (pH 7.5), 100 mM NaCl, 2 mM DTT. The sample was then extensively dialyzed against the same buffer to remove the DMSO using membranes of 0.5-1 kDa molecular weight cut-off. Crystals were obtained at room temperature by vapor diffusion mixing equal volumes of the β4-BP230 complex at 2.5 mg/ml and crystallization solution. The best crystals were obtained using as crystallization solution 0.1 M Na-acetate (pH 5.0), 20% PEG 6000, 0.2 M MgCl_2_. Crystals were transferred to a similar solution containing 25% glycerol and were flashed-cooled in liquid nitrogen. Data were collected at 100 K on the Xaloc beam line of the ALBA synchrotron (Barcelona, Spain) (Juanhuix et al., 2014). A high multiplicity dataset was obtained by combining three sets of 1800 images, 0.2° oscillation per image, measured at three positions of a single crystal. Diffraction data of these and all other crystals were processed with the XDS suite (Kabsch, 2010).

Crystals belong to the space group C2 (Table 1) and contain one copy of the complex in the asymmetric unit (∼46% solvent content). The structure was phased by molecular replacement with the program Phaser (McCoy et al., 2007) using the structures of the individual FnIII-3 and FnIII-4 domains (PDB codes 4WTW and 4WTX). The structure was refined against data extending to 1.55 Å resolution with phenix.refine (Afonine et al., 2012), alternated with model building in Coot (Emsley et al., 2010). Refinement included overall anisotropic and bulk solvent corrections, positional refinement, restrained refinement of individual B-factor, and refinement of the translation/libration/screw-rotation (TLS) parameters of eight groups in β4 and two in BP230. The refined model had excellent geometry and included residues 1457-1481, 1486-1552, and 1557-1665 of β4, residues 27-50 of BP230, 104 molecules of water, and two molecules of glycerol. Detailed statistics of the refinement are shown in Table 1.

Samples of the WT β4-FnIII-3,4 in complex with BP230 26-55, were prepared as for the mutant. Crystals were obtained using as crystallization solution 0.1 M phosphate-citrate (pH 4.2), 20% PEG 8000, 0.2 M NaCl. Crystals were cryoprotected with 25% glycerol in the crystallization solution and were flashed cooled in liquid nitrogen. Diffraction data were collected at 100 K on the i03 beam line of the Diamond Light Source (Didcot, United Kingdom).

Crystals of the WT β4-BP230 complex were isomorphous with the β4(T1663R)-BP230 crystals. The structure was solved starting from the coordinates of the mutant complex and was refined against data extending to 2.05 Å. After rigid body fitting, the refinement was done as for the mutant, with the exception that only four TLS groups were refined. The refined model included residues 1457-1483, 1486-1551, and 1561-1666 of β4, residues 27-49 of BP230, and 46 molecules of water (Table 1).

### SAXS measurements and analysis

SAXS data were measured at the P12 beamline of the European Molecular Biology Laboratory (EMBL) at the Deutsches Elektronen-Synchrotron (Hamburg, Germany) using radiation of wavelength (λ) 1.24 Å and a Pilatus 2M detector (Dectris) (Blanchet et al., 2015). Samples of β4-CS-FnIII-3,4 WT and point mutants were equilibrated in 20 mM Na-phosphate (pH 7.5), 150 mM NaCl, 5% glycerol, 3 mM DTT. Samples at various concentrations were prepared by 2-fold serial dilutions as indicated in the Table S1. SAXS data from protein samples and their buffers were measured consecutively at 10 °C. Data were collected for a range of the scattering-vector from 0.01 to 0.45 Å^−1^ (q=(4πsinθ)/λ, where *2θ* is the scattering angle). Data were processed and analyzed using the ATSAS package (Franke et al., 2017). Guinier analysis was done with the program AUTORG (Petoukhov et al., 2007). *P*(*r*) functions were calculated using data to *q* ≤ 0.30 Å^−1^ with the program GNOM (Svergun, 1992). Inter-domain flexibility was analyzed using the Ensemble Optimization Method with the program EOM 2.1 (Tria et al., 2015). First, a pool of 15.000 theoretical conformers that represent the potential conformation diversity of the β4-CS-FnIII-3,4 was generated treating the FnIII domains as rigid bodies and the CS and the linker as flexible regions. Then, combinations of the theoretical scattering profiles of the models were fitted to the experimental curves.

### Site-directed spin labeling and DEER measurements and analysis

The mutant proteins of β4-FnIII-3,4 containing two cysteines R1485C/C1608, R1504C/C1608, and L1497C/C1608, all of which also carried the changes C1559A/C1483S/T1663R, were labeled with the thiol-reactive paramagnetic probe S’-(1-oxyl-2,2,5,5-tetramethyl-2,5-dihydro-1^-pyrrol-3-yl)methyl methanesulfonothioate (MTSL) as described (Alonso-Garcia et al., 2015). Solutions of MTSL-labeled proteins between 70 to 100 ^M in 20 mM Na-phosphate (pH 7.5), 150 mM NaCl, without or with ∼5-fold molar excess of BP230 26-55, were mixed with deuterated glycerol in a 2:1 ratio to obtain a vitrified solution with longer transverse relaxation times upon freezing; they were subsequently transferred into 3 mm OD quartz tubes and stored in liquid nitrogen until measurement.

DEER measurements were performed at a temperature of 50 K in a Q-band (∼ 34 GHz) home-made EPR spectrometer (Gromov et al., 2001) equipped with a rectangular TE(102) cavity allowing for big sample volumes (Tschaggelar et al., 2009). The two-frequency 4-pulse DEER sequence (Pannier et al., 2000) was used setting the pump pulses at the maximum of the nitroxide spectrum. The DEER traces were processed with DeerAnalysis, an ad hoc toolbox programmed for MATLAB (Jeschke et al., 2006) and analyzed using a model-free Tikhonov regularization implemented in the same software to obtain the distance distributions.

### Structure and sequence analysis

Pairwise superimposition of structures was done using the program LSQKAB (Kabsch, 1976). Domain motions that relate the BP230-bound and the free structures of β4-FnIII-3,4 were analyzed with the program DynDom (Hayward and Berendsen, 1998). Evolutionary conservation scores were calculated using the Consurf Server (Ashkenazy et al., 2016). Prediction of the sequence tolerance within the β4-binding site of BP230 was done using the Sequence Tolerance method (Smith and Kortemme, 2011) as implemented in the ROSIE Server of the Rosetta molecular modeling package (Lyskov et al., 2013). Molecular figures were created with PyMOL (Schrödinger, 2015). Analysis of potentially phosphorylatable sites was done using the Phosphonet server (Kinexus Bioinformatics) (http://www.phosphonet.ca/).

### Antibodies

The following primary antibodies were used: mouse monoclonal antibody (mAb) 450-11A against β4 (BD Biosciences), human mAb 5E against BP230 (Ishiko et al., 1993), mouse mAb 233 against BP180 (Nishizawa et al., 1993), guinea pig polyclonal antibody (pAb) P1 against plectin (Stegh et al., 2000), and rabbit pAb D-8 against FLAG (Santa Cruz Biotechnology). Secondary antibodies were as follows: goat anti-mouse Texas Red (Invitrogen), goat anti-human Alexa Fluor 488 (Invitrogen), goat anti-guinea pig Alexa Fluor 488 (Invitrogen), and donkey anti-human Texas Red (Jackson ImmunoResearch).

### cDNA constructs for expression in mammalian cell cultures

Point mutations R1463E, R1463A, C1559A, and T1663R were introduced by site directed mutagenesis (see above) in a construct coding full-length β4 in the pUC18 vector (Niessen et al., 1997b). Retroviral vectors with mutant β4 cDNAs were generated by subcloning the mutant β4 cDNAs in the LZRS-MS-IRES-ZEO vector as described (Frijns et al., 2012).

The cDNA coding for the region 19-162 of BP230 was amplified by PCR using primers that added *EcoR*I and *Not*I sites at each end, respectively. The PCR product was digested and was cloned using the same sites in the pCEF-FLAG vector (Chiariello et al., 2000), which codes for an N-terminal FLAG tag. Point mutations were introduced in this construct by site-directed mutagenesis.

### Cell culture and immunofluorescence

Immortalized keratinocytes derived from a patient with Pyloric Atresia associated with Junctional Epidermolysis Bullosa (PA-JEB) have already been described (Schaapveld et al., 1998). PA-JEB/β4 keratinocytes stably expressing WT or point mutants of β4 were generated by retroviral transduction as described (Sterk et al., 2000). For immunostaining, PA-JEB/β4 cells were seeded on glass coverslips, fixed in 1% paraformaldehyde in PBS, permeabilized with 0.2% Triton X-100 in PBS for 5 min, blocked with 2% BSA in PBS, and incubated for 1 h with the following primary antibodies: 5E anti BP230 (dilution 1:500), 450-11A against β4 (dilution 1:500), P1 against plectin (dilution 1:200), and 233 against BP180 (undiluted). Cells were washed three times with PBS and were incubated with the secondary antibodies for 1 h. Cells were washed three times with PBS, stained with 4’,6-diamidino-2-phenylindole (DAPI, Sigma), and washed again three times with PBS. Finally, the coverslips were mounted onto glass slides in Mowiol-DABCO and observed using a Leica TCS SP5 confocal microscope.

The degree of co-localization between BP230 and β4, plectin or BP180 was quantified using the Manders’ overlap coefficients (MOC) calculated with the JACoP plugin from Image-J (Bolte and Cordelieres, 2006). Statistical significance was analysed using unpaired, nonparametric Mann-Whitney test in GraphPad.

### Pull-down assay

HEK293T cells were maintained in Dulbecco’s modified Eagle’s medium supplemented with 10% fetal bovine serum. Transfection with pCEF-FLAG-BP230 (19-162) constructs was done using polyethylenimine (PEI) (PEI:DNA ratio 2:1). 48 h after transfection, cells from a confluent 10-cm plate were lysed in 600 μl lysis buffer consisting of 20 mM Tris (pH 7.5), 500 mM NaCl, 0.5% Triton X-100, 1 mM Na_3_VO_4_, 25 mM NaF, 1 mM PMSF, and protease inhibitor cocktail (Roche). Lysates were cleared by centrifugation at 16000 x *g* for 15 min at 4 °C. Supernatants were incubated for 1 hour at 4 °C with 0.6 nmoles of the GST-β4 (1436-1666) T1663R or the inactive double mutant R1463E/T1663R and 20 μl of glutathione-agarose resin (Agarose Bead Technologies). The resin was washed four times with lysis buffer and the bound proteins were extracted with SDS-PAGE sample buffer. Bound FLAG-BP230 proteins were analyzed by immunobloting using anti-FLAG antibody (dilution 1:1000) and as secondary antibody a goat anti-rabbit IgG DyLight 800 (Thermo Fisher) (dilution 1:5000). Results were detected by infrared fluorescence using an Odyssey imaging system (Li-Cor).

### Data availability

Coordinates and structure factors have been deposited to the Protein Data Bank (PDB) under the accession numbers 6GVL (β4(WT)-BP230) and 6GVK (β4(T1663R)-BP230); raw diffraction images have been deposited in the Zenodo repository (DOI 10.5281/zenodo.1287191 and 10.5281/zenodo.1286853). SAXS data have been deposited in the Small Angle Scattering Biological Data Bank (SASBDB) under codes SASDDE8, SASDDF8, SASDDG8, SASDDH8, SASDDJ8, SASDDK8, SASDDL8, SASDDM8, SASDDN8, and SASDDP8. Other data are available from the corresponding authors upon reasonable request.

## Acknowledgments

We thank Prof Gunnar Jeschke for access to the EPR spectrometer. Lisa te Molder is thanked for assistance with image analysis and statistical analysis of data. We thank Constantina Bakolitsa for critical comments. We acknowledge ALBA-CELLS, Diamond Light Source (proposal MX10121), and the EMBL for access to synchrotron radiation facilities. This work was supported by the Spanish Ministry of Science, Innovation and Universities (grants BFU2009-08389, BFU2015-69499-P, and CTQ2015-64486-R) and Aragón local government by the action Grupo de Referenda “Biología Estructural”; these grants were co-funded by the European Regional Development Fund (ERDF). MGH, AC, and NAG were recipients of predoctoral training grants from Universidad de Salamanca, Spanish Ministry of Education, Culture and Sport (FPU14/06259), and Consejo Superior de Investigaciones Científicas, respectively. This research received funding from the European Community’s Seventh Framework Programme (FP7/2007-2013) under BioStruct-X (grant agreement N°283570). JMdP’s institution is supported by the Programa de Apoyo a Planes Estratégicos de Investigación de Estructuras de Investigación de Excelencia co-funded by Castilla-León autonomous government and ERDF (CLC-2017-01).

## Author contributions

JAM, IGR, AS, JMdP, Conception and design, Acquisition of data, Analysis and interpretation of data, Writing the article; MGH, AC, Conception and design, Acquisition of data, Analysis and interpretation of data; NAG, MK, Acquisition of data, Analysis and interpretation of data.

