## Supplemental figures S1 to S10 and table S1 for "Integrin α6β4 recognition of a linear motif of bullous pemphigoid antigen BP230 controls its recruitment to hemidesmosomes"

### SUPPLEMENTAL MATERIAL

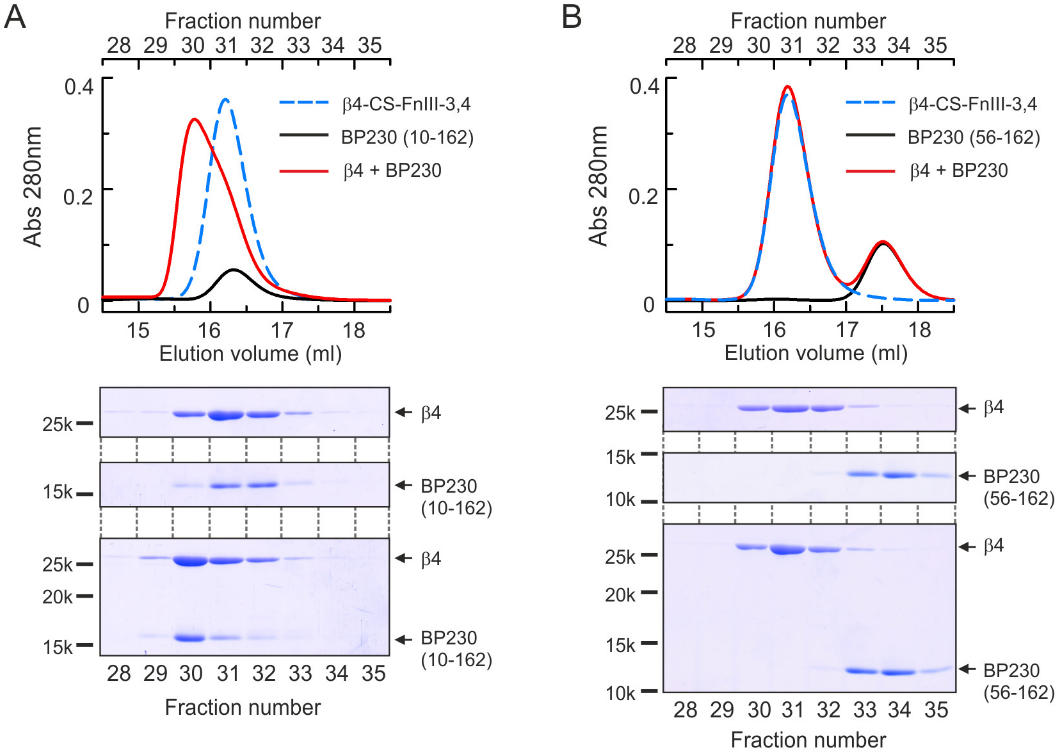


**Figure. S1. Analysis by size exclusion chromatography of the interaction between β4-CS-FnIII-3,4 and the BP230 fragments 10-162 (A) and 56-162 (B)**. Chromatograms of the isolated β4 and BP230, and equimolar mixtures are shown. Samples were analyzed using a Superdex 200 10/300 GL column (GE Healthcare) equilibrated in 20 mM Tris (pH 7.5), 150 mM NaCl, 0.5 mM tris(2-carboxyethyl)phosphine. The injection volume was 100 µl, the flow rate was 0.5 ml/min, and fractions of 0.5 ml were collected. Analyses of the fractions by SDS-PAGE are shown aligned in the bottom.

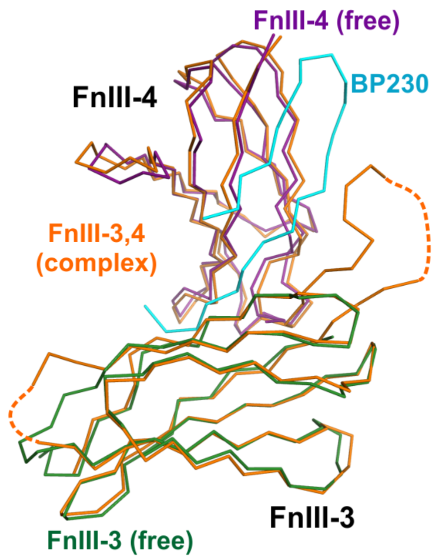


**Figure S2**. **Comparison of the FnIII domains in the isolated structures and in the β4-BP230 complex**. Superimposition of the individual structures of the FnIII-3 (dark green, PDB code 4WTW) and FnIII-4 (dark violet, PDB 4WTX) domains onto the equivalent regions of the β4-BP230 structure (β4 orange, BP230 cyan). The structures are shown as Cα traces.

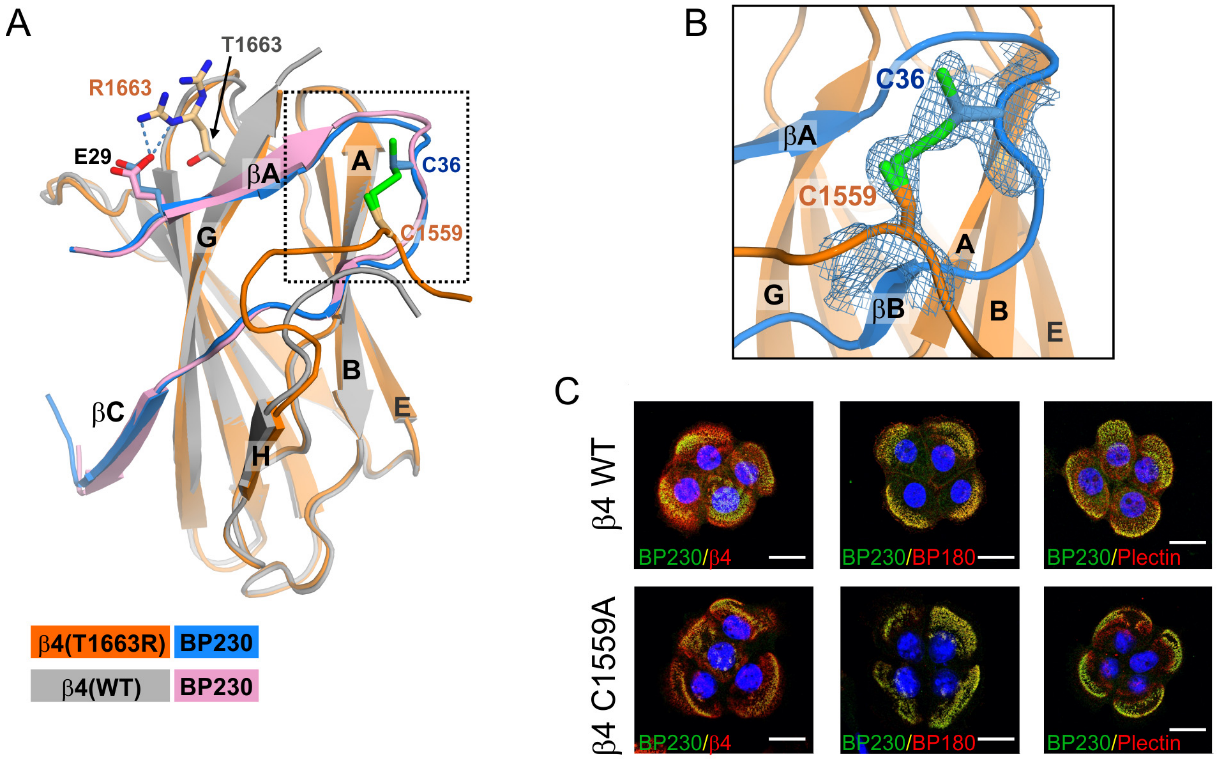


**Figure S3**. **Structure of the β4 inter-domain linker in the β4(T1663R)-BP230 complex**. **(A)** Ribbon representation of the β4(T1663R)-BP230 complex (β4 orange and BP230 blue) superimposed onto the β4(WT)-BP230 complex (β4 grey and BP230 violet). For clarity, the FnIII-3 is not shown. The side chains of β4 T1663 (WT), the mutant R1663, and E29 of BP230 are shown as sticks. The salt bridge between β4-T1663R and BP230-E29 is shown as dashed lines. The side chains of β4-C1559 and BP230-C36, which were modeled as a partially formed disulfide bridge in the β4(T1663R)-BP230 structure, are also shown. **(B)** Close-up view of the region around β4-C1559 and BP230-C36 in the β4(T1663R)-BP230 structure (dash square in A). A *2mFobs-DFcalc* map (contoured at 1 σ) is shown around the two cysteines. **(C)** Analysis of the role of the disulfide bond in the recruitment of BP230 in keratinocytes. Confocal microscopy images of PA-JEB keratinocytes expressing WT or C1559A β4. Cells were stained pairwise with antibodies against BP230 and either β4, BP180, or plectin. Nuclei were counterstained with DAPI (blue). Scale bar, 20 μm. When β4-C1559A was expressed in PA-JEB keratinocytes, it recruited endogenous BP230 to a similar extent as when WT β4 was expressed, suggesting that C1559 is not required for the interaction with BP230. Thus, a physiological role cannot be assigned to the disulfide bridge, which might have formed during the crystallization.

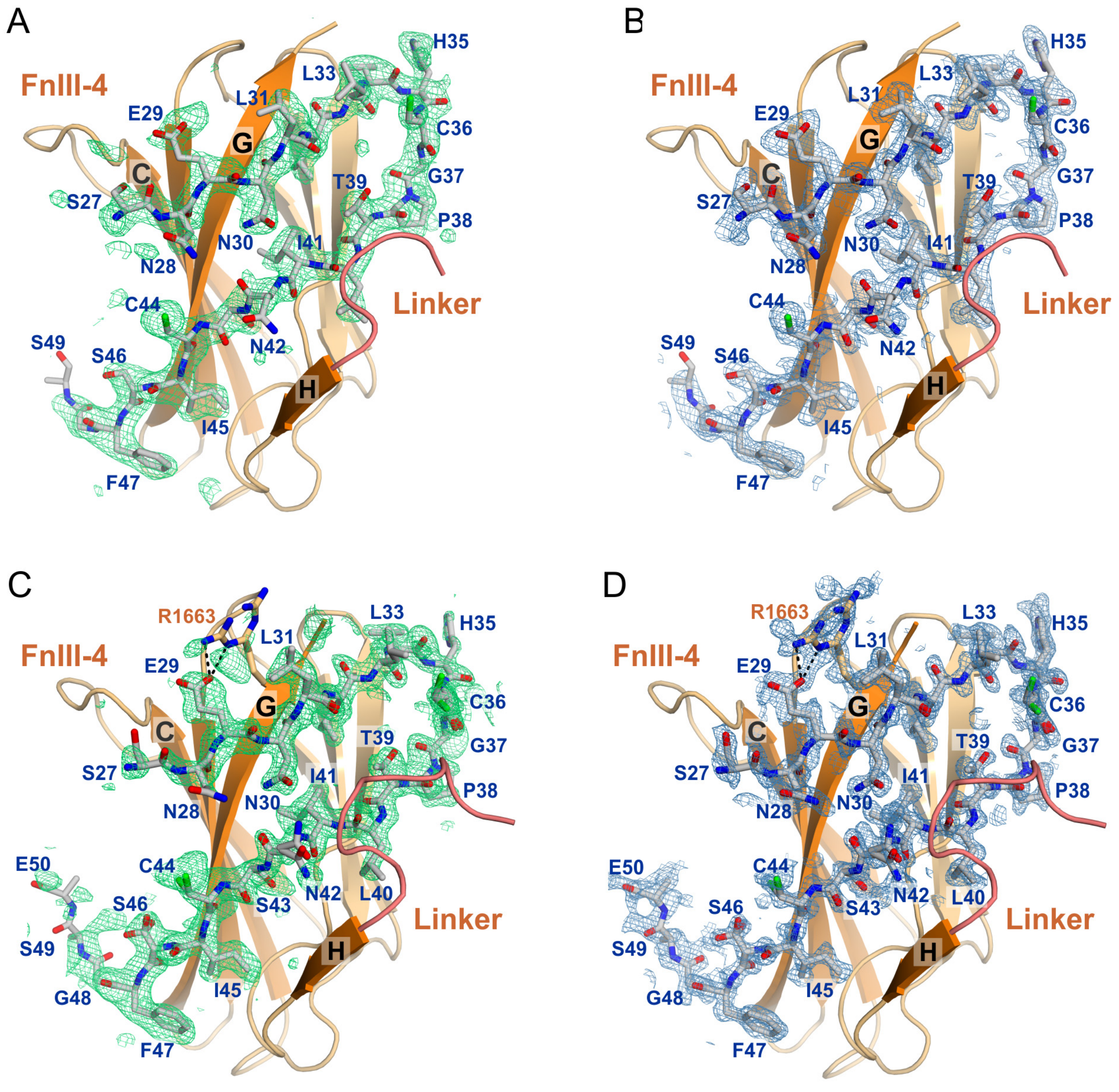


**Figure S4**. **Electron density for BP230 in the structures of β4(WT)-BP230 (A, B) and β4(T1663R)-BP230 (C, D)**. (**A, C**) Composite omit difference maps (*mFobs-DFcalc*) contoured at 2.5 σ around BP230 (shown as sticks). Model bias was reduced by using a simulated annealing refinement protocol and by sequentially omitting parts of BP230 from the refinement. (**B, D**) Feature-enhanced maps (FEM), *2mFobs-DFcalc*, contoured at 1.5 σ around the BP230. For clarity, the FnIII-3 domain is not shown.

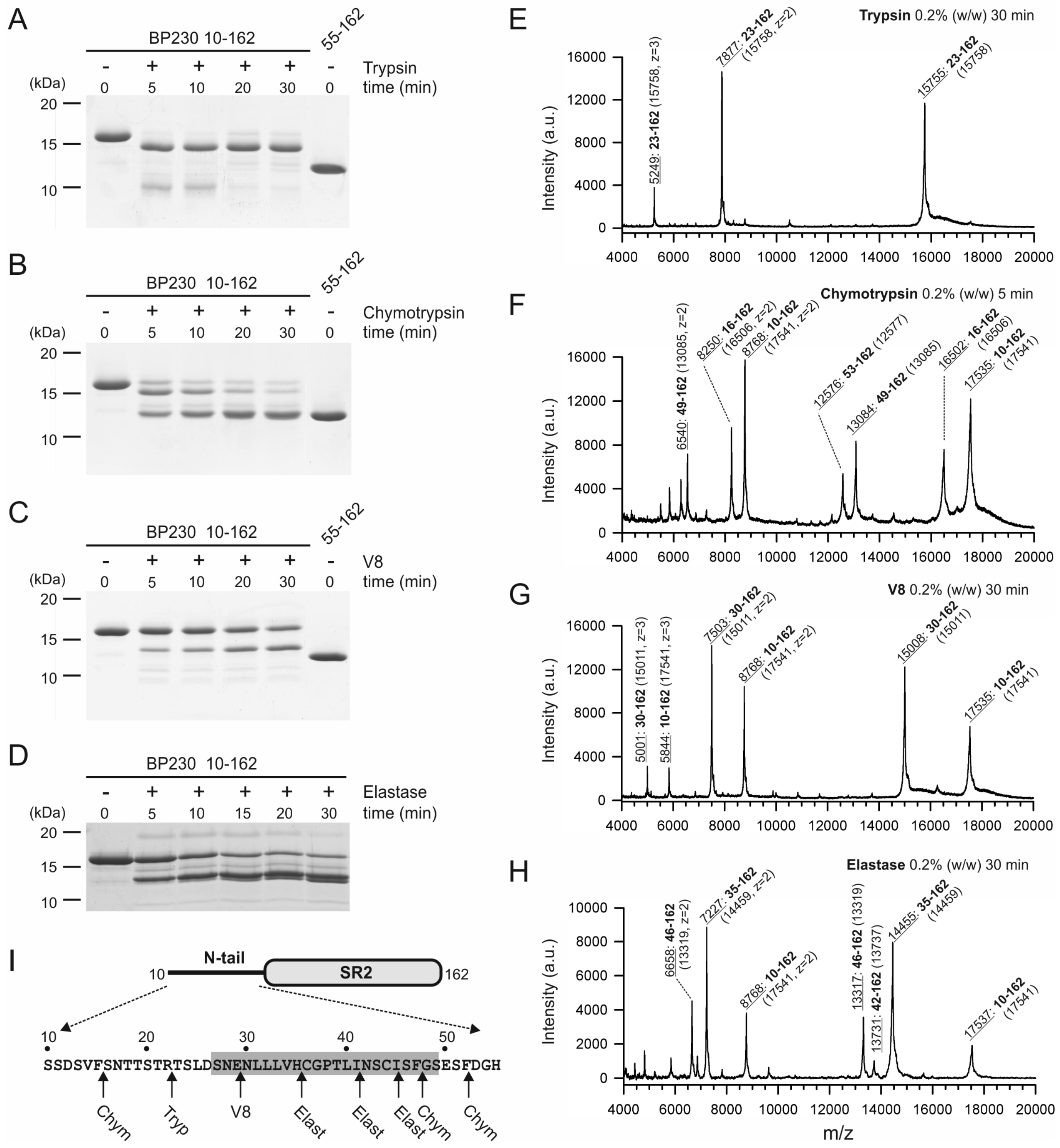


**Figure S5**. **Limited proteolysis of BP230 in the absence of β4**. (**A-D**) Analysis by SDS-PAGE (14% acrylamide gels) of time course digestions of BP230 10-162 with 0.2% (w/w) trypsin (A), chymotrypsin (B), protease V8 (C), or elastase (D). In A-C the last lane corresponds to the SR2 domain (55-162). (**E-H**) analysis by MALDI-TOF mass spectrometry of the masses of the proteolytic fragments of BP230 10-162 as indicated. The experimental mass of each major peak is underlined. The corresponding region of BP230 is shown in bold and the calculated masses are in parenthesis. (**I**) Representation of the domain structure of BP230 10-162. All the digestion sites are located in the N-tail as indicated under the sequence. The β4-binding region is highlighted by a grey box.

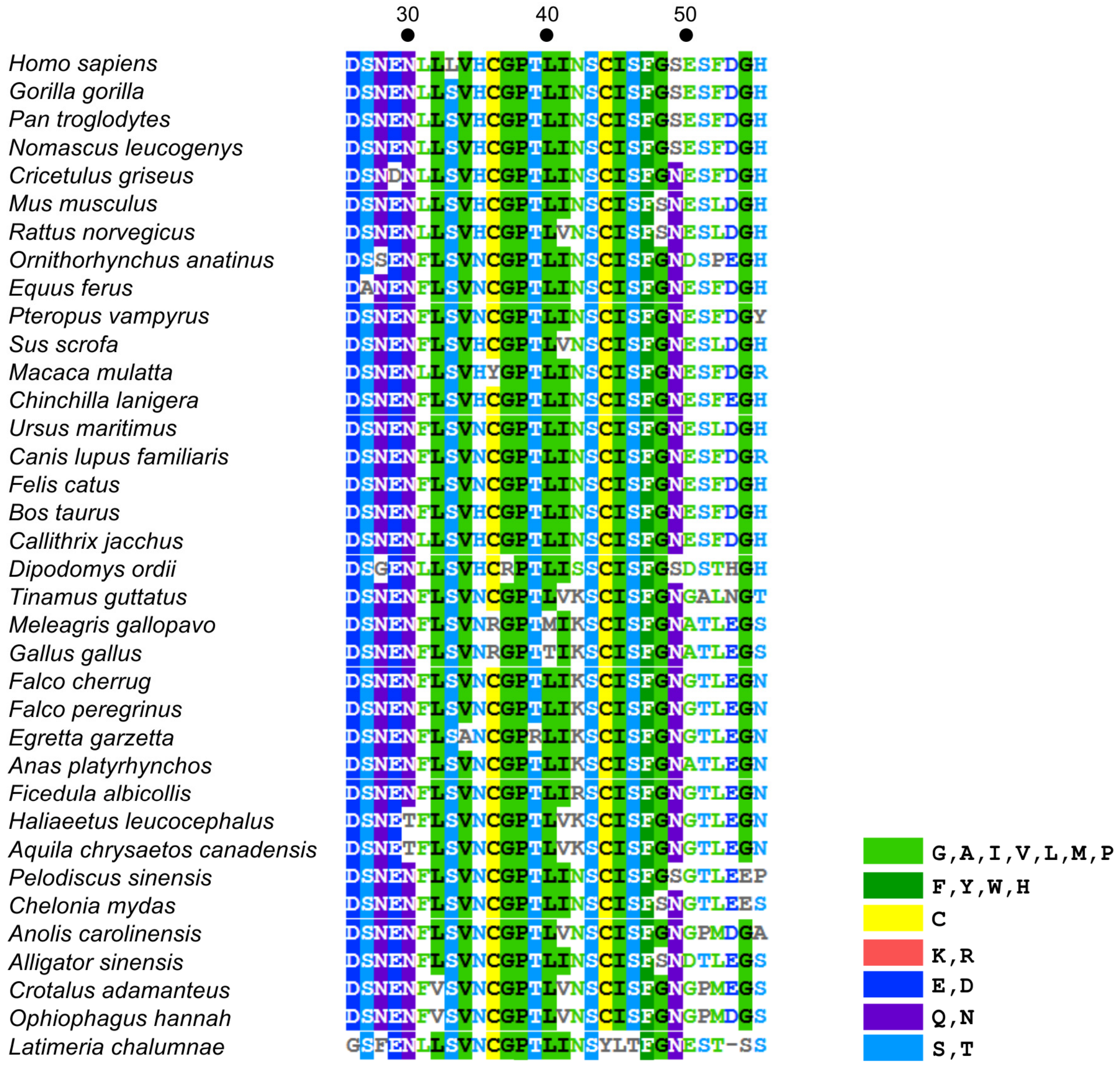


**Figure S6**. **Alignment of the β4-binding region of BP230 from multiple species**. The region 26-55 of human BP230 was aligned with the equivalent segments of orthologs from 35 species. Residues that match the consensus sequence (80%) are highlighted by boxes colored according to the type of residue as indicated in the legend.

**Predicted rank** (rows) for each BP230 position (columns); residue numbers 30, 35, 40, 45 are printed above N, H, L, I.

| BP230 | E | N (30) | L | L | L | V | H (35) | C | G | P | T | L (40) | I | N | S | C | I (45) | S | F | G |
|---|---|---|---|---|---|---|---|---|---|---|---|---|---|---|---|---|---|---|---|---|
| 1 | T | H | T | **L** | **L** | **V** | G | Q | **G** | **P** | I | R | T | L | A | K | **I** | **S** | **F** | **G** |
| 2 | V | **N** | R | Q | F | W | N | A | N | G | F | D | **I** | D | G | Q | V | A | W | S |
| 3 | K | Q | **L** | E | R | I | D | S | S | S | V | K | G | R | D | R | L | E | H | A |
| 4 | I | S | K | A | W | K | K | R | D | A | W | **L** | A | T | **S** | A | E | R | R | D |
| 5 | F | A | Y | S | Y | R | R | E | R | Q | R | M | V | E | T | S | A | G | S | N |
| 6 | R | D | F | M | I | Q | S | N | A | E | D | S | S | M | E | G | T | T | Y | T |
| 7 | Y | E | V | N | M | M | Q | K | H | K | A | A | D | S | N | E | D | D | A | Q |
| 8 | A | K | S | D | K | T | M | G | Q | R | L | N | N | A | V | D | M | N | V | R |
| 9 | S | M | A | T | E | A | A | T | W | M | H | Q | E | K | Q | M | S | I | L | E |
| 10 | Q | R | I | G | V | S | W | M | E | H | K | E | Q | **N** | I | N | N | Q | Q | P |
| 11 | **E** | W | M | K | Q | E | F | D | F | D | S | V | M | Q | L | T | G | V | D | L |
| 12 | M | G | Q | I | S | N | L | L | Y | Y | Q | T | H | G | K | W | Q | K | N | V |
| 13 | D | T | W | R | T | H | E | H | K | W | M | I | K | I | H | F | R | M | K | C |
| 14 | H | F | P | V | G | D | Y | F | M | N | **T** | G | L | V | M | Y | K | L | M | H |
| 15 | L | Y | H | H | D | L | **H** | V | L | L | E | H | R | P | P | L | H | H | G | K |
| 16 | W | V | E | F | H | G | T | I | T | T | N | W | F | H | R | H | Y | Y | E | I |
| 17 | N | I | G | Y | A | Y | V | Y | V | V | Y | F | Y | F | F | V | F | P | I | M |
| 18 | G | L | N | P | N | F | I | W | I | F | G | Y | W | C | Y | I | P | F | T | Y |
| 19 | P | P | D | C | P | P | P | P | P | I | P | P | P | Y | W | P | W | W | P | F |
| 20 | C | C | C | W | C | C | C | **C** | C | C | C | C | C | W | C | **C** | C | C | C | W |

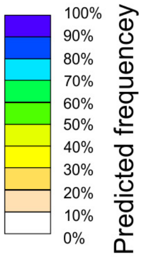


**Figure S7. Prediction of the sequence tolerance in the β4-binding site of BP230**. Table of the amino acids ranked individually for each sequence position of the region 29-48 of BP230 according to the frequency with which they are predicted to be tolerated without significantly affecting the stability of BP230 and the binding interface. The effects of the amino acid substitutions were predicted with Rosetta (using a Boltzmann factor *kT* = 0.59) using the structure of the WT β4-BP230 complex. The BP230 sequence is shown above. Wild type residues in the ranked list are shown in red.

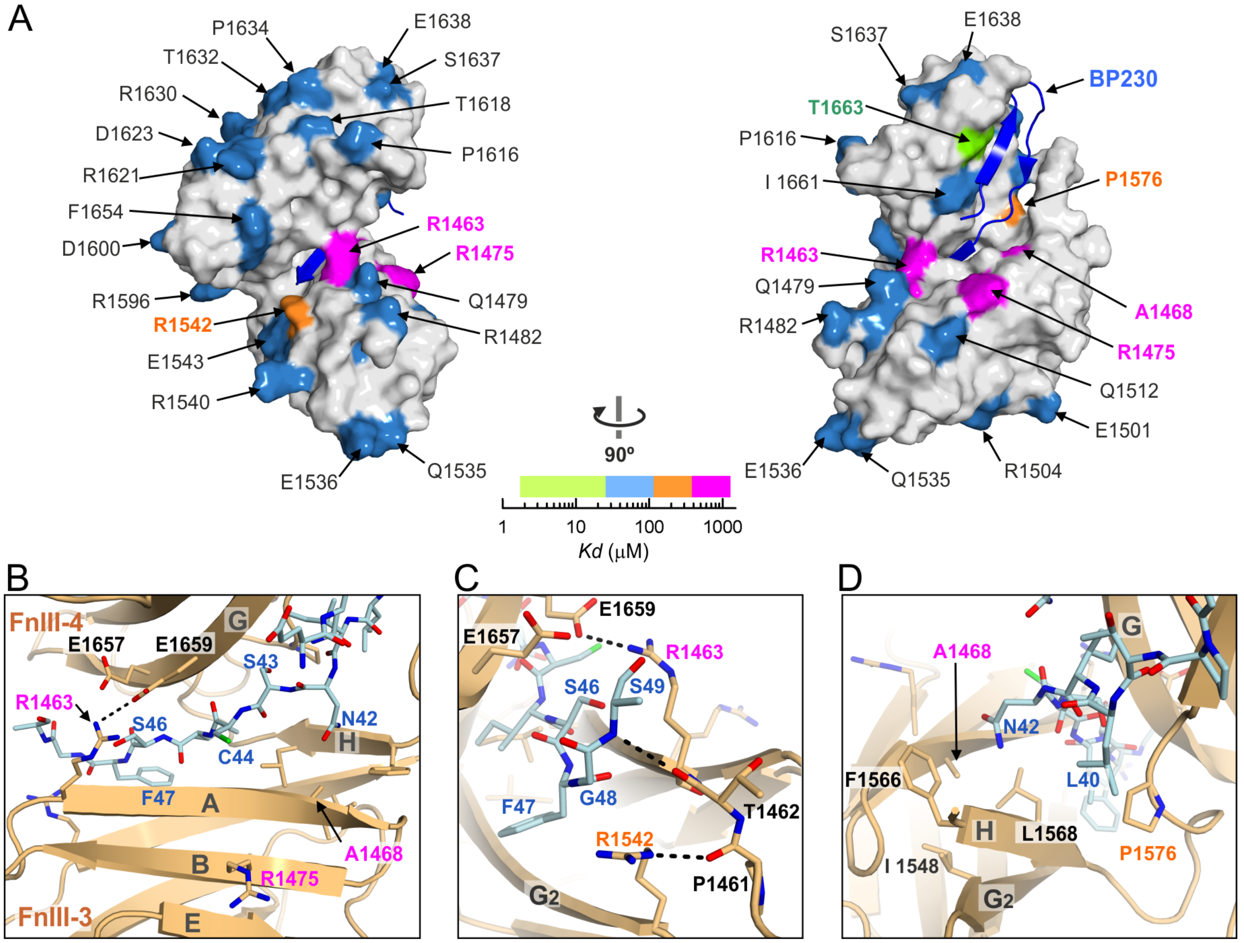


**Figure S8. Structural interpretation of the mutagenesis data**. (**A**) Structure of the β4(WT)-BP230 complex. BP230 is shown as a ribbon and β4 is shown in a surface representation. Residues are colored according to the effect of their mutations as in Fig 2. (**B-D**) Close ups of regions that include residues of β4 that were shown in the mutagenesis analysis to be important for binding of BP230.

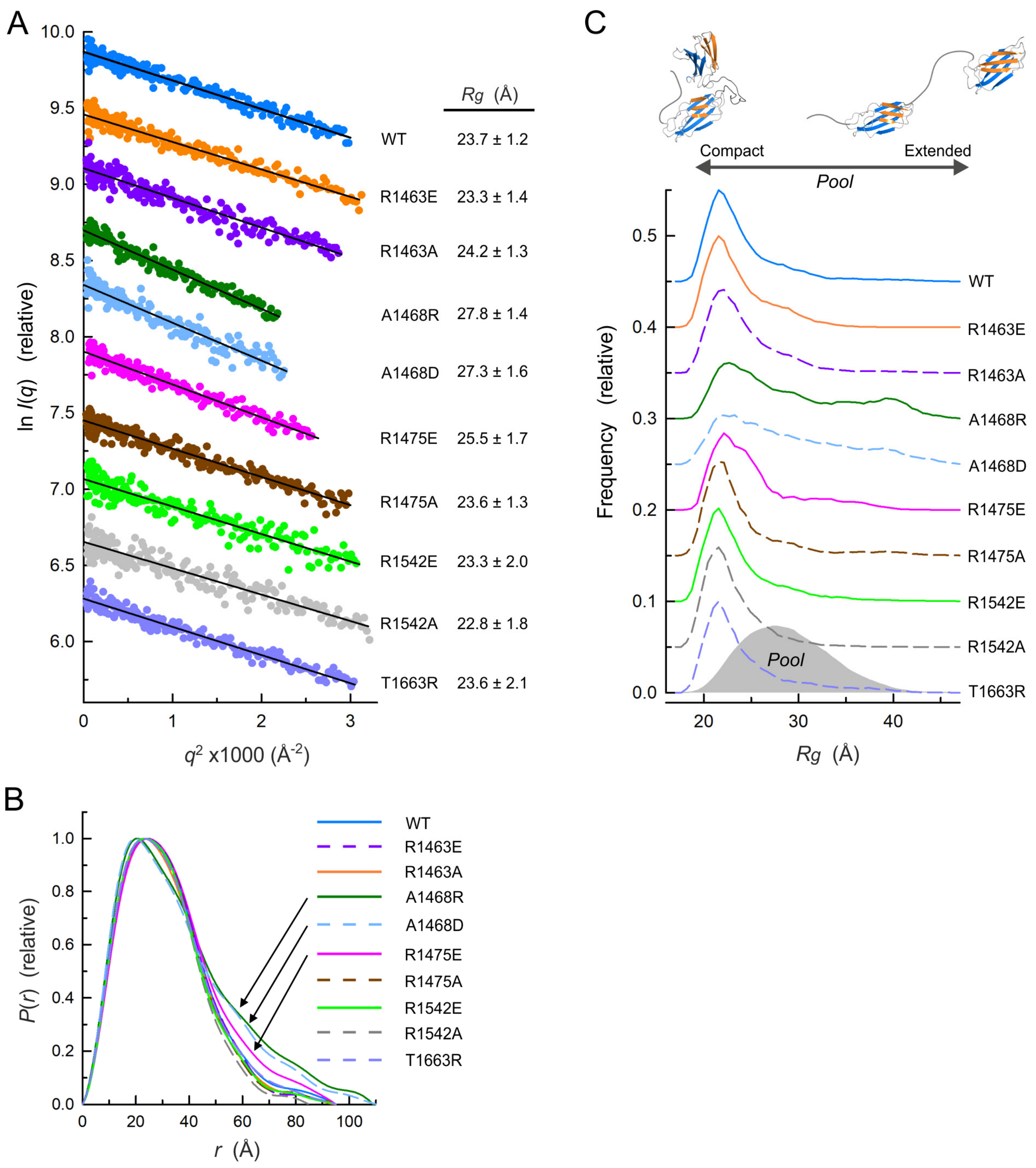


**Figure S9. Analysis by SAXS of the effect of point mutations on the structure of the CS-FnIII-3,4 of β4**. (**A**) Guinier plots of the SAXS data of β4-CS-FnIII-3,4 WT and mutants and the derived *Rg* values (±STD). Lines are the Guinier fit. (**B**) *P*(*r*) functions of β4 WT and mutants. (**C**) EOM analysis of the flexibility in the WT and mutants of β4-CS-FnIII-3,4. The frequency distributions of *Rg* in a pool of calculated models (grey area) and in the selected ensembles that fit the SAXS data of the WT and mutant proteins (lines) are shown. Plots in A and C are vertically displaced for representation.

**Table S1. SAXS data collection and derived parameters of β4-FnIII-3,4 proteins**

| | WT | R1463E | R1463A | A1468R | A1468D | R1475E | R1475A | R1542E | R1542A | T1663R |
|---|---|---|---|---|---|---|---|---|---|---|
| **Data collection** | | | | | | | | | | |
| Concentration range (mg $ml^{-1}$) | 0.9 – 14.9 | 1.4 - 5.5 | 0.9 - 7.1 | 1.5 - 11.9 | 0.9 - 14.9 | 1.2 – 9.2 | 1.3 – 10.5 | 0.9 – 14.8 | 0.9 – 14.5 | 1.5 – 11.7 |
| Exposure time (sec) | 30 x 0.05 | 30 x 0.05 | 30 x 0.05 | 30 x 0.05 | 30 x 0.05 | 30 x 0.05 | 30 x 0.05 | 30 x 0.05 | 30 x 0.05 | 30 x 0.05 |
| **Structural parameters** | | | | | | | | | | |
| Guinier analysis | | | | | | | | | | |
| $qR_g$ range | 0.15 - 1.30 | 0.15 - 1.30 | 0.18 - 1.30 | 0.16 – 1.30 | 0.16 – 1.30 | 0.20 – 1.29 | 0.13 – 1.30 | 0.15 – 1.29 | 0.15 – 1.30 | 0.13 – 1.30 |
| $I(0)/c$ ($10^{-2}$ $cm^2$ $mg^{-1}$) [a] | 1.60 | 1.58 | 1.64 | 1.62 | 1.53 | 1.89 | 1.43 | 1.54 | 1.52 | 1.55 |
| $R_g$ (Å) | 23.7 ± 1.2 | 23.3 ± 1.4 | 24.2 ± 1.3 | 27.8 ± 1.4 | 27.3 ± 1.6 | 25.5 ± 1.7 | 23.6 ± 1.3 | 23.3 ± 2.0 | 22.8 ± 1.8 | 23.6 ± 2.1 |
| $P(r)$ analysis | | | | | | | | | | |
| $I(0)/c$ ($10^{-2}$ $cm^2$ $mg^{-1}$) [a] | 1.62 | 1.59 | 1.64 | 1.64 | 1.53 | 1.90 | 1.44 | 1.55 | 1.51 | 1.57 |
| $R_g$ (Å) | 25.2 | 24.4 | 24.4 | 29.7 | 28.7 | 26.6 | 24.3 | 24.2 | 23.2 | 24.9 |
| $D_{max}$ (Å) | 95 | 95 | 95 | 109 | 110 | 95 | 95 | 95 | 85 | 95 |
| Porod volume, Vp ($Å^3$) | 36900 | 35400 | 36100 | 35300 | 34200 | 42300 | 35400 | 35100 | 34100 | 35300 |
| Molecular mass (kDa) [from (Vp/1.5)] | 24.6 | 23.6 | 24.1 | 23.5 | 22.8 | 28.2 | 23.6 | 23.4 | 22.7 | 23.5 |
| Monomeric mass from sequence (kDa) | 25.6 | 25.6 | 25.6 | 25.6 | 25.6 | 25.6 | 25.6 | 25.6 | 25.6 | 25.6 |
| SASBDB code | SASDDE8 | SASDDF8 | SASDDG8 | SASDDH8 | SASDDJ8 | SASDDK8 | SASDDL8 | SASDDM8 | SASDDN8 | SASDDP8 |

[a] Absolute intensities determined using water as a secondary standard

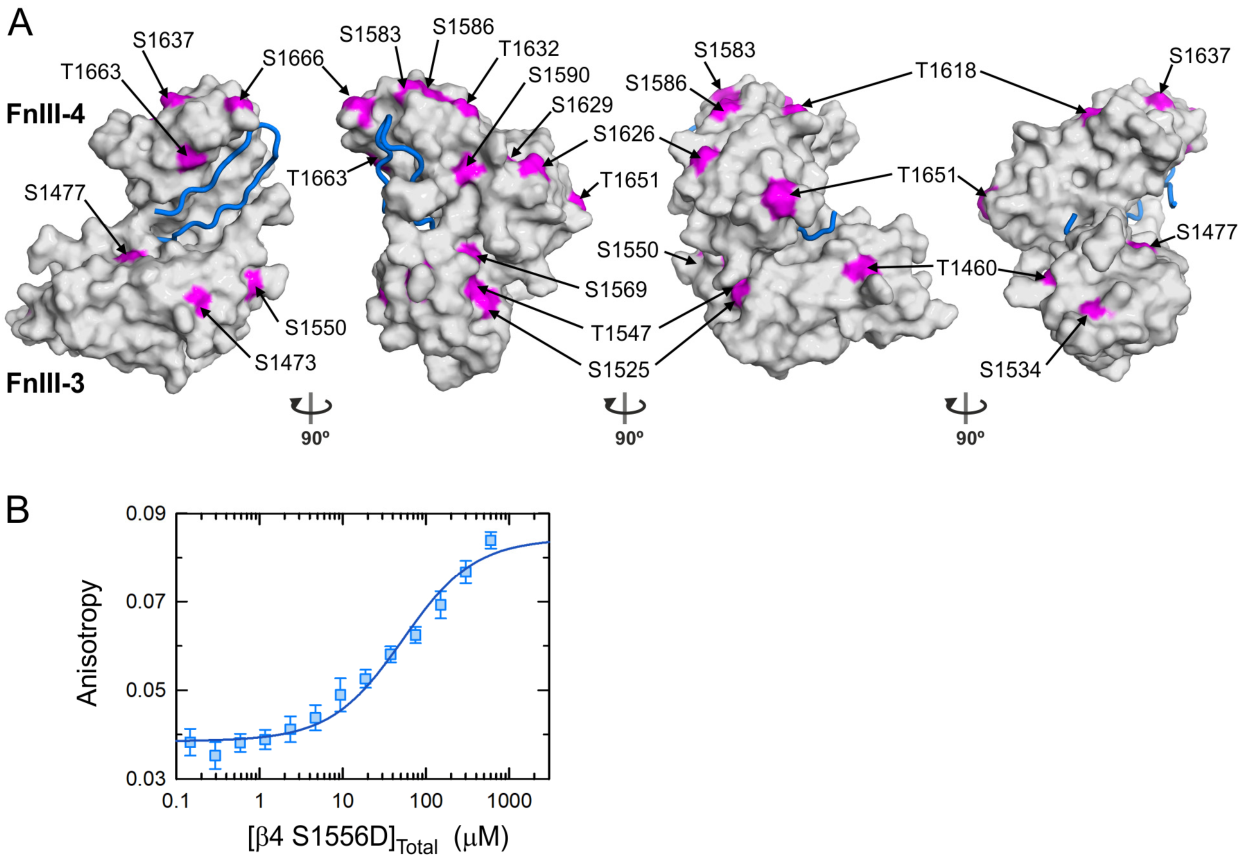


**Figure S10. Putative phosphorylatable serine and threonine residues in the FnIII-3,4 of β4**. (**A**) Surface representation of the FnIII-3,4, with the backbone of BP230 shown as a blue wire. Ser/Thr residues in β4 predicted to be potential phosphorylation sites by the Phosphonet server are shown in pink. (**B**) Binding of β4 (1457-1666) carrying the phosphomimetic substitution S1556D to the fluorescein-labeled BP230 peptide 26-55 (0.5μM), measured by fluorescence anisotropy. The line represents the fit to the data, which corresponds to *Kd* = 52 ± 11 μM (± standard error).
